# Ancestral sequence reconstruction enables functional expression of archaeal thermostable tungsten-containing aldehyde:ferredoxin oxidoreductases in *E. coli*

**DOI:** 10.64898/2026.11.16.767025

**Authors:** Deborah M. Boes, Yutong Wang, Puck de Wit, Rob A. Schmitz, Javier Viña, Ivan Mateljak, Frank Hollmann, Peter-Leon Hagedoorn

## Abstract

Tungsten-containing enzymes offer important opportunities in industrial biocatalysis, e.g. to convert carboxylic acids into aldehydes without requiring co-substrates ATP and NADPH. These W-enzymes operate at low reduction potentials compared to the more ubiquitous Mo-enzymes, enabling unique reactions. Their application is limited by the lack of functional expression in commonly used heterologous host organisms. Here, we show that variants of *Pyrococcus furiosus* aldehyde:ferredoxin oxidoreductase (*Pf*AOR) obtained using ancestral sequence reconstruction can be functionally expressed in *E. coli* with an activity of 0.86 U/mg towards crotonaldehyde at 60°C, which is a 50-fold higher activity compared to the recombinant extant enzyme. The ancestral enzymes, named *Thermo*AOR3 and *Thermo*AOR4, carrying 61 and 84 mutations respectively, were purified using a single-step affinity purification, showed a *k_cat_* of 0.95 and 0.86 s^-1^ at 60 °C, respectively, and have *K*_m_ values comparable to native *Pf*AOR. *Thermo*AOR3 contained a redox active [4Fe-4S] cluster like native *Pf*AOR. Furthermore, it contained 10-15% of the expected pterin cofactor, W and Fe, which indicates the enzyme production can still be improved. Altogether, we demonstrate the first heterologous expression of a functionally active W-AOR using a widely used bacterial host, which represents a breakthrough enabling exploration of these enzymes for industrial biocatalysis.

## Introduction

Metalloenzymes are biocatalysts containing at least one metal cofactor to mediate catalysis. Since most transition metals can adopt multiple valence states, they can mediate electron transfer and facilitate redox reactions. Of all transition elements, tungsten (W) is the heaviest element known to be used in biological systems ^1^. In enzymes, the W ion is incorporated in a W-cofactor consisting of a metal center coordinated by two ligands, called metallopterin (MPT) or pyranopterin dithiolene ^2,3^.

In some enzymes, molybdenum (Mo) and W are interchangeable as metal cofactor due to their similar chemical properties. This replacement changes the properties of the cofactor due to the difference in reduction potential of the two metals ^4^. Tungsten complexes generally exhibit lower reduction potentials, often by 200 – 300 mV, compared to the corresponding molybdenum-containing complexes ^5^. The effect of this metal substitution was demonstrated in the dimethylsulfoxide reductase (DMSOR) from *Rhodobacter capsulates.* Substitution of the naturally occurring Mo by W lowered the reduction potential of the cofactor by approximately 220 – 330 mV ^6^.

Tungsten (W)-dependent enzymes offer a largely untapped repertoire of catalysts for organic synthesis. Their W-cofactors support transformations which are challenging in established synthetic chemistry. The unique low reduction potential enables W-enzymes to catalyze thermodynamically challenging redox reactions. The selective reduction of carboxylic acids to aldehydes represents a particularly compelling opportunity for this chemistry (Fig. 1). Carboxylic acids are abundant starting materials, whereas aldehydes provide versatile intermediates for alcohol and amine synthesis and carbon-carbon bond formation. Yet their interconversion requires reduction of a poorly reactive carboxyl group while suppressing further reduction of the more reactive aldehyde and preserving other functional groups. Selective chemical methods have been developed, including manganese- and nickel-catalyzed routes, but representative systems rely on stoichiometric silanes, substrate activation or subsequent release of a protected aldehyde ^7,8^. ATP-dependent carboxylic acid reductases (CARs) provide a biocatalytic solution by coupling substrate adenylylation to NADPH-dependent reduction ^9^. Their requirement for both ATP and reducing cofactors asks for alternative catalytic strategies that bypass ATP-dependent substrate activation. W-dependent aldehyde oxidoreductases (AORs) offer such an alternative. *Pyrococcus furiosus* (*P. furiosus*) aldehyde: ferredoxin oxidoreductase (AOR), naturally oxidizes aldehydes to carboxylic acids using its physiological redox partner ferredoxin, but is also able to catalyze the thermodynamically challenging reverse reaction, the reduction of non-activated carboxylic acids to aldehydes (*E’_m_* ≈ −560 mV), if low-potential electron donors are available ^10–12^. It contains a W center coordinated by two pyranopterin dithiolene ligands, forming a W-bis-MPT cofactor, together with a [4Fe-4S] cluster for electron transfer. Besides AOR, other AOR-family enzymes have been purified and characterized from the native archaeal host *P. furiosus*, including glyceraldeyde-3-phosphate:ferredoxin oxidoreductase (GAPOR), formaldehyde oxidoreductase (FOR), WOR4 and WOR5. Some of these enzymes are highly specific towards one substrate, such as GAPOR for D-glyceraldehyde-3-phosphate, while AOR shows substrate promiscuity and catalyzes the reversible oxidation of a range of different aldehydes ^13–16^. Native purification and recombinant production in hosts such as *Aromatoleum evansii* and *Thermoanaerobacter kivui* have already enabled AOR characterization, but large-scale production is yet to be established ^17,18^.

**Figure 1.**
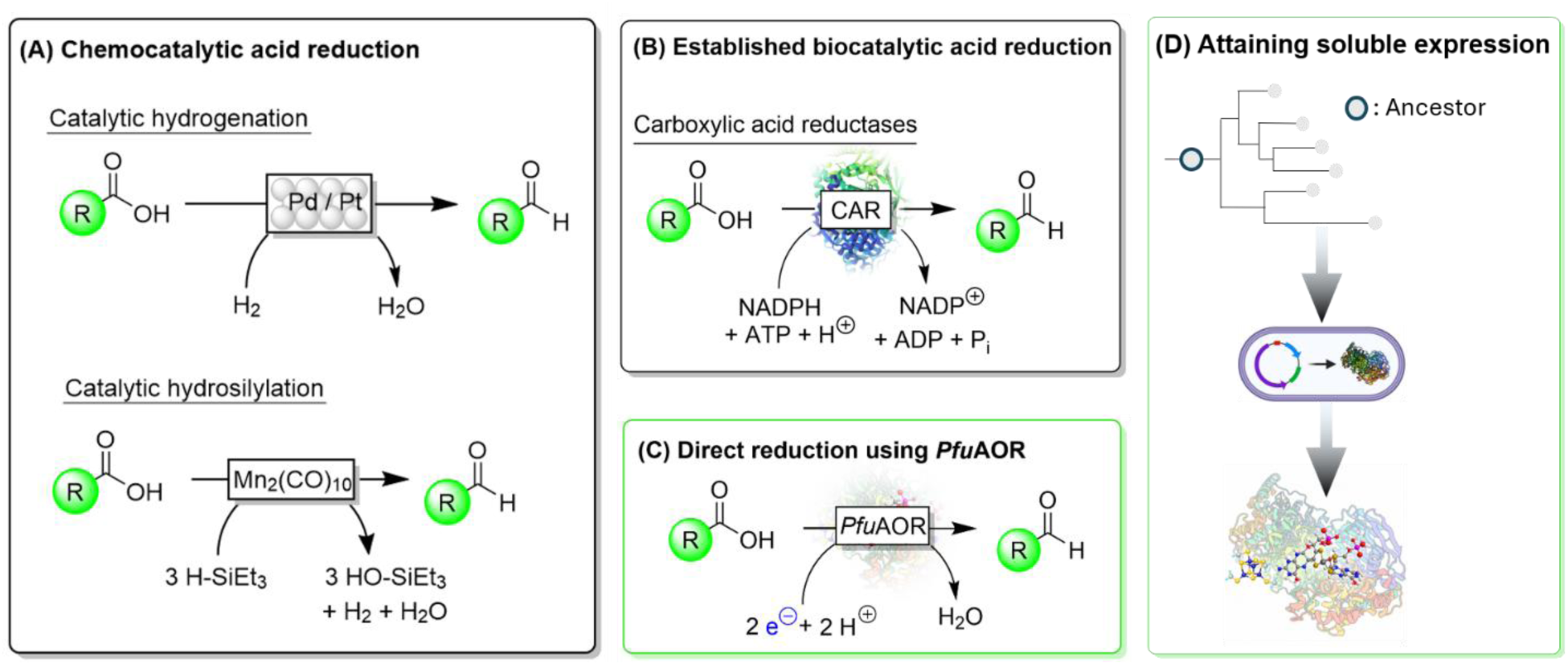
W-AORs offer an alternative biocatalytic solution for the conversion of carboxylic acids to aldehydes. **a** chemocatalytic carboxylic acid reduction, **b** established biocatalytic acid reduction, **c** W-AOR biocatalytic acid reduction, **d** enabling W-AOR biocatalytic applications by functional expression in *E. coli*.

Recently, other W-containing enzymes have gained interest as promising biocatalysts for capturing and recycling the green-house gas CO_2_. A W-dependent formate dehydrogenase (FDH) catalyzes CO_2_ reduction and produces formate, a C1 chemical feedstock ^19^. FDH contains a different W-cofactor than AOR, namely a W-bis-MGD (molybdopterin guanine dinucleotide) cofactor, and is part of a different enzyme family than AOR. In comparison to the W-bis-MPT cofactor found in AOR-family enzymes, each pterin ligand includes guanosine monophosphate (GMP) to form the W-bis-MGD cofactor ^20^. Using the potential of W-enzymes requires reliable access to active enzymes. For that reason, considerable research is focused on the production of these FDHs in heterologous hosts to obtain them on a larger scale, because the actual application of FDHs, AORs, and other W-containing enzymes, such as DMSORs, is held back by limitations in their production ^19,21^.

The challenge, however, extends beyond soluble expression to assembly and incorporation of the W-bis-MPT cofactor and iron–sulfur cluster, ideally in an expression system that also allows for genetic engineering. The most widely used host for heterologous expression of enzymes, the bacterium *Escherichia coli*, does not naturally produce W-enzymes and most heterologously produced W-enzymes may therefore end up as inactive apo-enzymes or non-functional inclusion bodies ^22^. It has to be noted that most of these investigated W-enzymes, such as FDHs and DMSORs, can exist either as Mo- or W-containing enzymes that have a similar structure and amino acid sequence ^23^. *E. coli* encodes an endogenous Mo-FDH and may therefore be more amenable to produce a W-FDH. In contrast, AOR family enzymes exclusively exist as W-containing enzymes, making heterologous production in an organism that does not naturally produce these W-enzymes even more challenging. Interestingly, the *E. coli* genome contains an AOR family homolog ydhV. The ydhV gene is not expressed under any known conditions but has been characterized after recombinant production ^24^. However, no activity was found for this protein.

To improve heterologous expression of a stable and active archaeal protein in a bacterial host, we used ancestral sequence reconstruction (ASR) to predict ancestral variants of AOR. ASR is increasingly being used as a protein engineering technique, because it has been shown to produce more robust proteins with enhanced stability and solubility ^25^. Moreover, previous studies have shown that predicted ancestral enzymes can exhibit an altered or broader substrate scope compared to the extant enzymes ^26^. Indeed, Kajimoto et al. (2022) showed that the production of an ancestral RNA ligase resulted in increased thermostability and activity and even appeared to enhance protein crystallization for structure determination compared to the extant counterpart ^27–29^.

In this study, we analyzed if ancestral sequence reconstruction could improve the production of functional W-containing AOR enzymes in *E. coli*. By designing and expressing proteins belonging to different nodes of a phylogenetic tree based on *P. furiosus* AOR (*Pf*AOR), we characterized three evolutionary ancestors of *Pf*AOR, named *Thermo*AOR3, *Thermo*AOR4, and *Thermo*AOR5. Our studies revealed that *Thermo*AOR3 and *Thermo*AOR4 have an up to 50-fold higher specific activity compared to wild-type *Pf*AOR heterologously expressed in the same *E. coli* strain under optimized cultivation conditions. This specific activity represents circa 5% of the activity of native *Pf*AOR isolated from *P. furiosus*. *Thermo*AOR3 showed an incorporation of the MPT-ligand, Fe, and W to 10-20% of the maximal theoretical values, indicating that the obtained activities can still be enhanced when cofactor incorporation is improved. The *Thermo*AORs were produced at a level of 7 ± 0.7 mg per L culture and isolated using a single-step affinity chromatography purification, making the production process scalable for future industrial applications.

## Materials and Methods

### Ancestral sequence reconstruction

A comprehensive dataset of W-AOR sequences based on *Pyrococcus furiosus* W-AOR was assembled using the Protein BLAST (https://blast.ncbi.nlm.nih.gov/Blast.cgi) tool from the National Center for Biotechnology Information (NCBI). Redundant, truncated, or poorly annotated sequences were manually removed.

A multiple sequence alignment was generated using the MUSCLE algorithm implemented in the MEGA12 (Molecular Evolutionary Genetics Analysis) software ^30^. Alignment curation involved removal of outlier sequences and gaps, trimming of poorly aligned regions, and ensuring conservation of catalytic residues.

To construct phylogeny, the Bayesian method implemented in the program MrBayes 3.2.7a was used with 44 extant sequences as input ^31^. This analysis used two independent Markov chain Monte Carlo runs, with 16 chains each, and was performed for 10,000,000 generations to ensure convergence. Trees were sampled every 1000 generations for each run, resulting in a total of 20,000 trees, from which a consensus tree was obtained with high probabilities at the nodes. Selected ancestral nodes were reconstructed using PAML4 version 4.9j with the Wag substitution model and incorporating the gamma distribution for variable replacement rates across sites ^32^.

For each node, a single consensus ancestral sequence was generated, with very high (> 0.9) posterior probabilities guaranteeing robust reconstruction confidence. Profiles of probability for the most probable residue versus position of the sequences for the reconstructed proteins *Thermo*AOR3, *Thermo*AOR4 and *Thermo*AOR5 are given in Fig. S1.

### Expression of *Pf*AOR and ASR proteins in *Escherichia coli*

For the production of the ancestral sequence reconstructed (ASR) proteins in *E. coli*, plasmids were created containing the codon-optimized protein sequences of the proteins. The codon-optimized genes were synthesized by Synbio Technologies (Table S1, New Jersey, USA). Subsequently, the gene constructs were placed in multiple cloning site 1 (MCS-1) of a pETDuet-1 vector in the same reading frame as an N-terminal His-tag using restriction enzyme digestion with BamHI and HindIII (Fig. S2, Fig. S3, Fig. S4). In the pETDuet-1 plasmid, the genes are under the control of a T7 promotor to allow for controlled protein expression upon induction with isopropyl ß-D-thiogalactopyranoside (IPTG).

To produce wild-type *Pyrococcus furiosus* AOR recombinantly, the gene encoding AOR (locus PF0346, UniProt ID Q51739) was codon-optimized and synthesized by Synbio Technologies (Table S1, New Jersey, USA) in a pUC19 vector. Subsequently, the gene insert was cut from the pUC19 vector and placed in a pTrcHis A vector using restriction enzyme digestion with NcoI and HindIII. The genes are inducible using IPTG to allow for a controlled expression.

The wild-type *P. furiosus* AOR was expressed in multiple *E. coli* strains to study the difference in expression and protein production. These strains include *E. coli* MC1061, BL21(DE3), C43(DE3), and Rosetta-Gami 2(DE3, Sigma-Aldrich, St. Louis, Missouri, USA). The ASR proteins *Thermo*AOR3, *Thermo*AOR4, and *Thermo*AOR5 were only expressed in *E. coli* Origami 2(DE3).

To start the enzyme production, all plasmids were transformed into chemically competent *E. coli* cells by mixing 1 µL of plasmid (∼100 ng/µL) with 25 µL of cells. The cells were incubated on ice for 30 min after which they received a heat-shock at 42°C for 30 sec. After 2 min incubation on ice, 500 µL of Luria-Bertani (LB) medium was added and the cell suspension was incubated at 37°C and 400 RPM for 1 h to allow for expression of the antibiotic resistance gene. After incubation, 150 µL of cell suspension was plated on LB-agar plates containing 100 µg/mL ampicillin (Merck, Darmstadt, Germany) to select for the successfully transformed cells. L-Leucine and tetracycline were added to plates to cultivate *E. coli* Origami 2(DE3) or Rosetta-Gami 2(DE3) cells to final concentrations of 40 mg/L and 12.5 µg/mL, respectively. These cells have a Δ(ara-leu) mutation and therefore require additional leucine for growth. Next to that, they contain mutations in both the thioredoxin reductase (trxB) and glutathione reductase (gor) genes, which are selectable using a tetracycline selection marker.

After overnight incubation, a colony is picked to inoculate 10 mL LB medium containing 40 mg/L L-Leucine (only for Origami 2(DE3) and Rosetta-Gami 2(DE3) cells), and 100 µg/mL ampicillin. This pre-culture was incubated overnight at 37°C and 180 RPM. To start the overexpression of the proteins, 200 mL of Terrific Broth (TB) medium in 500 mL shake flasks (12 g/L bacto-tryptone (Becton, Dickinson and Company, Sparks, MD, USA), 24 g/L bacto-yeast extract (Becton, Dickinson and Company, Sparks, MD, USA), 4 mL/L glycerol (Merck, Darmstadt, Germany), 0.017 M KH_2_PO_4_, and 0.072 M K_2_HPO_4_ (Avantor, Radnor, PA, USA)) supplemented to create final concentrations of 5 g/L D-glucose (Merck, Darmstadt, Germany), 100 µg/mL ampicillin, 40 mg/L L-Leucine (for Origami 2(DE3) and Rosetta-Gami 2(DE3) cells), and 100 µg/mL Fe(III)NH_4_ citrate (Merck, Darmstadt, Germany) was inoculated with 1% pre-culture. The flasks were incubated at 37 °C and 180 RPM until the cells reached an optical density at 600 nm (OD_600_) of approximately 0.8. After reaching this OD_600_, the medium was supplemented with Na_2_WO_4_ (Merck, Darmstadt, Germany) to a final concentration of 1 mM. For the expression of the ASR proteins a final concentration 1 mM L-Cysteine (Merck, Darmstadt, Germany) was added during induction as well, to provide a sulfur source for incorporation of the [4Fe-4S]-cluster. Next, protein expression was induced by adding isopropyl β-D-1-thiogalactopyranoside (IPTG, CliniSciences, Nanterre, France) to a final concentration of 100 µM, unless otherwise specified. The cultures were incubated stationery for another 16 h at 37 °C, to gradually deprive the cells of oxygen and allow for a semi-anaerobic induction.

After induction, the cells were harvested by centrifugation at 7000 x *g* for 20 min at 4 °C (Sorvall RC6+, Thermo Fisher Scientific, Waltham, MA, USA), washed with 30 mL pre-cooled 50 mM KPi buffer pH 8.0, and centrifuged again at 7000 x *g* for 20 min at 4 °C to collect cell pellets.

### Purification of recombinant *Pf*AOR, *Thermo*AOR3, *Thermo*AOR4 and *Thermo*AOR5

From the purification onwards, all activities were conducted under strictly anaerobic conditions in a Coy anaerobic chamber (O_2_ < 2 ppm). Prior to use, all buffers were sparged with nitrogen gas for 45 min and placed open in the glovebox (M. Braun, Garching, Germany) at least a day before use to allow residual oxygen to diffuse out of the solution.

Cell pellets were transferred into the glovebox and resuspended in lysis buffer (4 mL/g cells, 50 mM KPi buffer pH 8.0, 1 mM phenylmethylsulfonyl fluoride (PMSF), 100 mM NaCl, 2 mM MgCl_2_, 1X FastBreak Reagent (Merck, Darmstadt, Germany), 0.1 mg/mL pancreatic DNase (Merck, Darmstadt, Germany), and 0.2 mg/mL lysozyme (Carl Roth GmbH C Co. KG, Karlsruhe, Germany)) and lysed in a thermoshaker (Eppendorf, Hamburg, Germany) at 25°C at 600 RPM for 30 min. Subsequently, the lysis mixture was centrifuged at 6000 RPM for 30 min in a tabletop centrifuge (Fisherbrand Clinal Centrifuge, Thermo Fisher Scientific, Waltham, MA, USA) and the cell-free extract (CFE) was collected.

Wild-type *Pf*AOR was purified using heat-purification by heating 500 µL of CFE at 80°C in a water bath for 30 min. During this heat treatment, all non-thermostable *E. coli* proteins precipitate, while the thermostable *Pf*AOR remains soluble. The precipitated proteins were removed by centrifugation at 13,000 RPM for 5 min in a tabletop centrifuge (Eppendorf, Hamburg, Germany).

His-tagged ASR AORs were purified using 1 mL His GraviTrap TALON gravity flow columns (Cytiva, Marlborough, MA, USA). Prior to loading the sample, the columns were equilibrated with 15 mL equilibration buffer (50 mM KPi buffer pH 8.0, 2 mM MgCl_2_, 10% (v/v) glycerol, 150 mM NaCl, 20 mM imidazole (pH 8.0)) and the CFE was mixed 1:1 (v:v) with equilibration buffer. The diluted supernatant was loaded onto the column and washed with 15 mL equilibration buffer. The bound proteins were eluted from the column using 3 mL elution buffer (50 mM KPi buffer pH 8.0, 2 mM MgCl_2_, 10% (v/v) glycerol, 150 mM NaCl, 300 mM imidazole (pH 8.0)). The imidazole was removed from the samples using a PD-10 desalting column (Cytiva, Marlborough, MA, USA). Purified protein fractions were collected and concentrated to approximately 200 µL using spin filters (Amicon Ultra Centrifugal Filter, 3 kDa MWCO). Protein concentrations were determined using a bicinchoninic acid (BCA) assay as described previously (Interchim, Montluçon, France) ^33^. Proteins were separated by SDS-PAGE using precast Criterion TGX Stain-Free Protein gels (Bio-Rad Laboraties Inc., Hercules, CA, USA).

### Aldehyde oxidation activity assay

To assess the activity of AOR, activity assays were performed under anaerobic conditions in a Coy anaerobic chamber (O_2_ < 2 ppm) at 60°C in 2 mL PMMA cuvettes with a path length of 10 mm. The aldehyde oxidation activity of AOR was routinely analyzed using crotonaldehyde as substrate and benzyl viologen (BV) as electron acceptor. The substrate conversion was indirectly measured by monitoring the reduction of BV at 600 nm (ε_600_ = 7,400 M^-1^ cm^-1^) ^34^. For the reaction, BV was added to EPPS buffer (50 mM, pH 8.4 at 60°C) pre-heated to 60°C to create a final concentration of 3 mM. Purified enzyme (5 µL, 2-4 mg/mL) was then added, and the background activity was measured on an Avantes Varius spectrophotometer (Avantes, Apeldoorn, The Netherlands). After stabilization of the background, the reaction was initiated with the addition of 0.5 mM crotonaldehyde (25 µL of a 40 mM stock, Avantor, Radnor, PA, USA) and the response in activity was monitored. The specific activity of the enzymes was calculated by subtracting the background activity from the activity after substrate addition and is visualized as micromoles aldehyde oxidized per minute per milligram protein (U/mg) under the assay conditions.

To analyze the substrate scope of *Thermo*AOR3, various substrates have been tested using the same benzyl viologen coupled activity assay as described above. These substrates include linear aliphatic, branched chain aliphatic, aromatic, and other aldehydes. All used substrates including, if necessary, the co-solvent used, can be seen in Table S3.

All used substrates were tested at a final concentration of 0.5 mM in the assay by adding 25 µL of a 40 mM stock solution to a final volume of 2000 µL, diluting the DMSO solvent, if used, to a final concentration of 1.25% (v/v) in the assay.

### Carboxylic acid reduction activity assay

To determine the acid-reduction activity of purified *Thermo*AOR3, hexamethyl viologen (HMV) was used as electron donor. The HMV was synthesized as described previously ^35^. During the reaction, the decrease in absorbance of reduced HMV at 595 nm (ε_595_ = 10,100 M^-1^ cm^-1^) was followed to determine the enzyme activity. First, 150 µL of an anoxic 6 mM HMV solution in KPi buffer (100 mM, pH 6.0 at 60°C) was pre-reduced using 4 µL 100 mM Ti(III)citrate. Next, 10 µL 175 mM phenylhydrazine was added to the reaction to trap the produced aldehyde after conversion. Another 4 µL 100 mM Ti(III)citrate solution was added to reduce HMV to an A_595_ of approximately 0.4. Subsequently, purified enzyme (2 µL, 2-4 mg/mL), followed by 5 µL of 2.1 M sodium acetate (Avantor, Radnor, PA, USA) were added and the oxidation of HMV was monitored. The final volume of the reactions was 175 µL. The specific activity of the enzymes was calculated by subtracting the background activity from the activity after substrate addition and is visualized as micromoles acid reduced per minute per milligram protein (U/mg).

### Gas chromatography analysis

Aldehyde conversion by *Thermo*AOR3 was quantified using GC analysis. Crotonaldehyde was used as a substrate with varying concentrations of 0, 0.56, 1.11, and 2.78 mM. Reactions were performed in glass GC vials in an anoxic environment in the glovebox. Final volume of the reaction was adjusted with anoxic EPPS buffer (50 mM, pH 8.4 at 60°C) to 300 µL. To all reactions, 15 µL 240 mM BV, 8 µL 3.55 mg/mL purified enzyme, and final concentrations of 0 – 2.8 mM crotonaldehyde were added. The reactions were incubated at 60°C for 4h after which they were transferred on ice outside of the glovebox. The reactions were stopped with the addition of 20 µL of 5 M HCl after which they were extracted twice with 150 µL EtOAc containing 5 mM n-dodecane (Acros Organics, Geel, Belgium) as internal standard. After extraction, the combined organic phase was analyzed by GC on a Shimadzu GC-2010 Plus system equipped with an FID-2010 Plus detector (Shimadzu Corporation, Kyoto, Japan). For detection a CP-Wax 52 CB column (Agilent Technologies, Santa Clara, CA, USA) was used. GC conditions and retention times can be seen in Table S4.

### Pterin quantification

To analyze the pterin content of the W-bis-MPT cofactors, a fluorometric assay was performed following procedures described previously ^18,36,37^. A 50 µL purified protein sample (∼2 mg/mL) was mixed with 200 µL of 55 mM KMnO_4_ in 0.1 M NaOH. This sample was heated at 95°C for 45 min in a water bath to oxidize the pterin cofactor into a fluorescent derivative, after which 700 µL ice-cold ethanol was added to precipitate the residual KMnO_4_. The precipitates were removed by centrifugation at 13,000 RPM for 15 min in a tabletop centrifuge (Eppendorf, Hamburg, Germany). The supernatant was directly analyzed in a fluorescence plate-reader (BioTek Synergy 2, Agilent Technologies, Santa Clara, CA, USA), with an excitation wavelength (λ_ex_) of 360 ± 40 nm and an emission wavelength (λ_em_) of 460 ± 40 nm using black 96 well plates (Greiner AG, Kremsmünster, Austria).

The pterin content in the samples was calculated by comparison with a calibration curve made with pterin-6-carboxylic acid (Merck, Darmstadt, Germany) as reference. For confirmation of the pterin presence, xanthine oxidase (Sigma-Aldrich, St. Louis, Missouri, USA) containing a Mo-MPT cofactor per monomer was used as positive control. All calibration and positive control samples were treated identical to the protein samples described above.

### Metal determination using ICP-MS

For the quantification of tungsten, iron, and molybdenum, samples were analyzed using inductively coupled plasma mass spectrometry (ICP-MS). All samples and standards were analyzed on a PerkinElmer NexION 2000 ICP-MS (nebulizer: glass, injector: glass) and were measured with a sensitivity starting from 0.1 ppb. Before analysis, protein samples were pre-treated by digestion with nitric acid. For the pre-treatment, purified protein sample (100 µL, 1-2 mg/mL) was mixed with 50 µL of 65% HNO3 and subsequently diluted with 350 µL of H_2_O. This sample was sonicated in an ultrasonic bath (Elmasonic S30, Elma Schmidbauer GmbH, Singen, Germany) for 10 min, followed by centrifugation at 13,500 RPM for 5 min in a tabletop centrifuge (Eppendorf, Hamburg, Germany). After centrifugation, 400 µL supernatant was mixed with 3600 µL of 2% HNO_3_ + 2% HCl to get to a final 50 times dilution of the protein sample. To check the quality of the measurements, samples with known concentrations of the analyzed elements ranging from 0 to 50 µM were prepared and treated using the same pre-treatment steps as described above. The following isotopes were used for quantification of the metals: ^56^Fe, ^98^Mo and ^184^W.

### UV-Vis spectroscopy

All UV-Vis spectroscopic measurements were performed on a Cary 60 spectrophotometer from Agilent Technologies at 21°C. The spectra were measured with 80 µL 3.5 mg/mL *Thermo*AOR3 in anoxic EPPS buffer (50 mM, pH 8.4 at 60°C) in a 1 mL quartz cuvette. The samples were added to the quartz cuvette as anoxic solutions in the Coy chamber (O_2_ < 2 ppm) and the cuvette was closed with a rubber stopper before being transported out of the glovebox. To observe the reduction of the [4Fe-4S]-cluster, sodium dithionite (final concentration 250 µM) was added. Spectra were recorded at low speed in a wavelength range from 200 nm to 800 nm using automatic baseline subtraction.

### EPR spectroscopy

EPR spectra were recorded on a Bruker EMXplus spectrometer with a helium-flow cryostat at 20 K, using the following EPR parameters: microwave frequency 9.406 GHz, microwave power 20 mW, a modulation frequency of 100 kHz, and a modulation amplitude of 10 G ^38,39^. In each experiment, protein sample (200 μL, circa 100 μM) was sealed in an EPR tube in a Coy anaerobic chamber (O_2_ < 2 ppm) followed by immediate freezing by liquid nitrogen.

### Protein structure prediction using Chai-1

The structures of the ancestral sequence reconstructed proteins were predicted using Chai-1 (Chai Discovery, San Francisco, CA, USA) ^40^. For the prediction, the amino acid sequences listed in supplementary Table S2 were used and the following SMILES (Simplified Molecular Input Line Entry System) notations for the ligands (Table 1). The Chai-1 predicted structures of the *Thermo*AORs are given in the supplementary information.

**Table 1.** SMILES notations used for protein structure prediction using Chai-1. The number of copies used refers to the amount per monomer.

| Ligand | Copies | SMILES notation |
|---|---|---|
| MPT cofactor | 2 | O=C1NC(N)=NC2=C1NC3C(OC(COP([O-])([O-])=O)C(S)=C3S)N2 |
| W atom | 1 | O=[W]=O |
| [4Fe-4S]-cluster | 1 | [S]12[Fe]3[S]4[Fe]1[S]5[Fe]2[S]3[Fe]45 |
| Crotonaldehyde | 1 | C/C([H])=C([H])/C([H])=O |

The active sites, as well as the protein surface, entry channels and surface charge have been visualized using the PyMol Molecular Graphics System, Version 3.1 (Schrödinger, LLC).

## Results and Discussion

### Expression of wild-type *Pf*AOR in various *E. coli* strains yields soluble thermostable enzyme

Recombinant expression in the well-established host organism *E. coli* is a common approach to obtain large protein quantities. However, using such a host organism to express W-enzymes is not trivial since both the [4Fe-4S] and W-bis-MPT cofactor need to be formed and incorporated properly. To assess whether *E. coli* can produce at least a minimal amount of functional W-enzyme, the gene encoding the wild-type protein of *P. furiosus* AOR was expressed in 4 different *E. coli* strains (Fig. S5). These strains were selected for initial screening to investigate whether their differences in cellular characteristics influence recombinant protein expression. These included K-12 derived strains (Rosetta-Gami 2(DE3), MC1061) of which Rosetta-Gami 2(DE3) also supplies rare tRNAs for infrequently used codons and BL21 derived strains (BL21(DE3), C43(DE3) ^41^. MC1061 has previously been successfully used for heterologous expression of the Mo-containing FDH FdsC from *Rhodobacter capsulatus* and may therefore be able to produce a similar metal-containing cofactor ^42^. Similarly, the Rosetta-Gami 2(DE3) strain has been used by Park et al. (2007) in research involving a Mo-dependent GAPOR enzyme ^43^. To date, the *Methanococcus maripaludis* GAPOR investigated by Park and coworkers is the sole reported example of a mesophilic GAPOR ^43^. As mentioned previously, all other known AOR-family enzymes are described as W-containing proteins. GAPOR, being a member of the AOR-family enzymes, contains a similar MPT cofactor. To assess the use of this bacterial strain in producing a W-containing enzyme of the same family, it was selected as one of the *E. coli* strains in the experiments. Rosetta-Gami 2(DE3) contains mutations in genes encoding glutathione and thioredoxin reductases. Both thioredoxin and glutathione can reduce disulfide bonds. The inactivation of these reductases keeps the proteins in an oxidized state which can aid the formation of disulfide bonds and can therefore enhance the expression of proteins that require disulfide bonds for proper folding ^44,45^.

After expression, the cells were harvested and lysed and the enzyme was purified using heat purification at 80 °C. During this heat purification all non-thermostable proteins, including most of the endogenous *E. coli* proteins precipitated while the thermostable *Pf*AOR remained in solution as shown by the dominant protein band at around 67 kDa, which corresponds to the size of the *Pf*AOR monomer (Fig. S5) ^46,47^.

Hence, *Pf*AOR can be induced and expressed as a soluble thermostable protein in *E. coli*. The amount of produced protein varied per host strain used for overexpression. Both strains visualized in Fig. S5 contain the same expression vector, but the amount of soluble AOR produced using *E. coli* Rosetta-Gami 2(DE3) is significantly higher than using MC1061 (Fig. S5a).

### *Pf*AOR can be expressed in *E. coli* with low but measurable activity and W-incorporation

Since successful recombinant protein production does not necessarily point to a properly functioning protein, all AORs produced in different *E. coli* strains were heat purified and further characterized. To start, we determined whether the produced enzymes show activity towards the oxidation of aldehydes. The AORs produced in *E. coli* strains MC1061, BL21(DE3) and C43(DE3) did not show any measurable activity towards crotonaldehyde, while AOR produced in Rosetta-Gami 2(DE3) had a specific activity of 17 ± 2 mU/mg (Fig. 2a).

**Figure 2.**
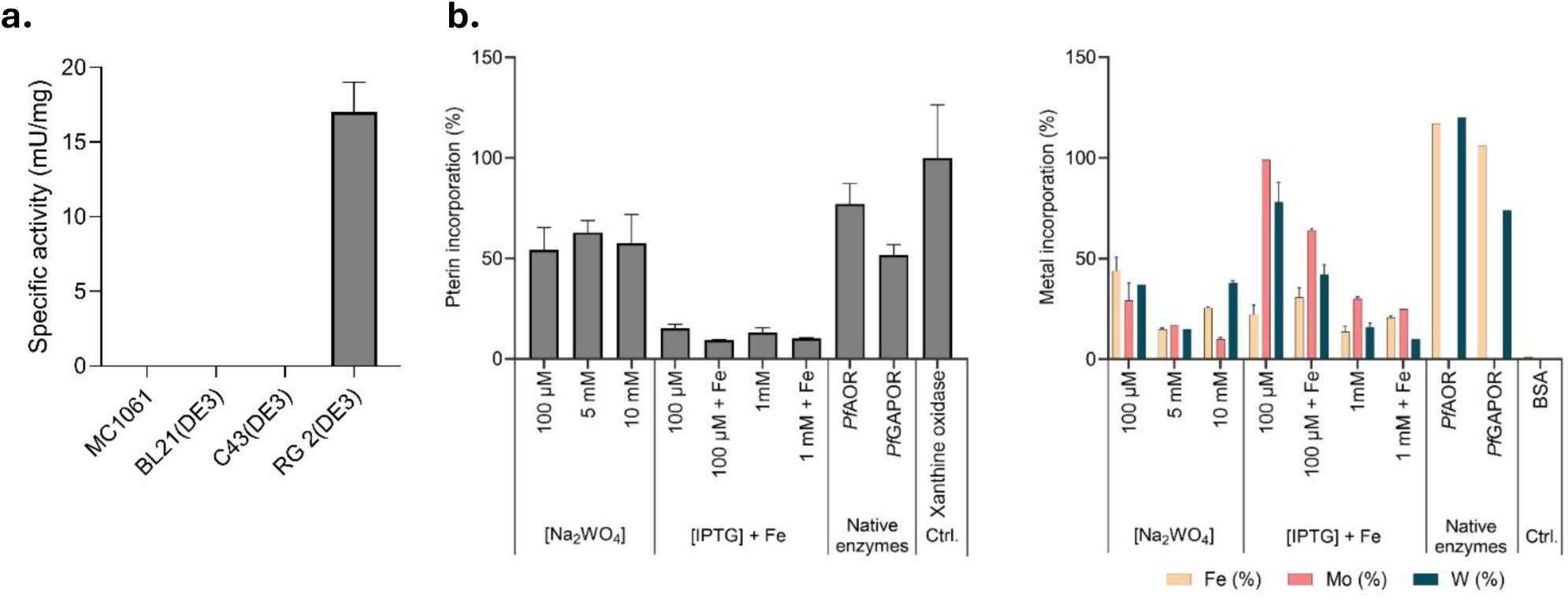
Activity and cofactor content of recombinant wild-type *P. furiosus* AOR. **a.** Oxidation activity of recombinant *Pf*AOR with crotonaldehyde as substrate. Activity was measured at 60 °C in an anoxic environment as crotonaldehyde-dependent reduction of BV (ɛ_600_ = 7,400 M^-1^cm^-1^). The reaction contained 3 mM BV, 0.5 mM crotonaldehyde and 0.4 – 1.5 mg AOR in EPPS buffer (50 mM, pH 8.4 at 60 °C). **b**. Pterin (right) and metal content (left) present in the W-bis-MPT cofactor and [4Fe-4S]-cluster. Changes in incorporated pterin and metals were measured upon addition of different concentrations of medium additives (Na_2_WO_4_, IPTG, and Fe(III)NH4 citrate (Fe)). The recombinantly produced proteins are compared to native AOR and GAPOR purified from *P. furiosus* cells. Left, Pterin content measured using a fluorescence assay with λ_ex_ of 360 nm and λ_em_ of 460 nm. The values are normalized to a pterin content of xanthine oxidase of 1 mol/monomer. Right, Molybdenum (Mo), Iron (Fe), and tungsten (W) content analyzed by ICP-MS. Bovine serum albumin (BSA) served as negative control. All values are visualized as percentage of incorporation relative to the expected values. Molybdenum is considered to have a maximum incorporation of 1 mol/monomer, equal to tungsten. All experiments were performed in duplicate, except the measurement of metal incorporation in native *Pfu* enzymes; in that case it was done once. [Na_2_WO_4_] – concentration sodium tungstate. [IPTG] – concentration IPTG. Ctrl. – Control. RG 2(DE3) – Rosetta-Gami 2(DE3).

Compared to the activity obtained with native AOR purified from *P. furiosus* cells, 27 U/mg at 65 °C, approximately 0.1% of the native activity could be obtained ^12^. It is worth noting that the enzyme activity is highly temperature dependent, with an approximate doubling of activity per 10°C when the temperatures fall within a range where exponential increase is observed ^12^.

Recombinant *Pf*AOR does not contain many cysteine residues (six in total), of which four are required to coordinate the [4Fe-4S] cluster. We therefore believe that the stability of disulfide bonds is not the major advantage of using Rosetta-Gami 2(DE3) for AOR production, but rather the slower growth observed when using Origami strains in general. The altered oxidized cytoplasm puts some physiological stress on the cells, which demands extra energy to maintain, compromising the growth rate ^48^. This slowed down growth, probably positively affected the folding of the proteins and incorporation of the cofactors ^49^.

The active *Pf*AOR obtained from the Rosetta-Gami 2(DE3) overexpression was further characterized to determine if both the [4Fe-4S]-cluster and W-bis-MPT cofactor were properly incorporated (Fig. 2b). The absolute values of incorporated metals and pterin cofactor can be seen in Fig. S6.

To simultaneously investigate whether the production of active enzyme was limited by the availability of one of the building blocks for the cofactor production, the effect of supplementing different concentrations of sodium tungstate (Na_2_WO_4_), IPTG and Fe(III)NH_4_ citrate (Fe) to the medium during protein expression was tested. When increasing the W concentration, the IPTG concentration was fixed at 100 µM and, similarly, when changing the IPTG concentration, the W concentration was fixed at 100 µM.

Increasing the Na_2_WO_4_ concentration in the medium from 100 µM to 10 mM caused no significant increase in incorporated tungsten (Fig. 2b, 37 and 38%, respectively) and pterin (Fig. 2b, 54 and 57%, respectively). Since *E. coli* lacks specific tungstate transporters, the tungstate is probably predominantly taken up in the cell via passive sulfate transport systems. The concentration gradient passively drives tungstate uptake into the cell. The lack of an increased W-incorporation when increasing the availability of sodium tungstate might indicate that the import of the metal is not the limiting factor for the production of the W-cofactor, but rather the lack of a specific cofactor biosynthesis pathway ^50^.

Molybdenum and tungsten share similar chemical properties, such as a similar ionic radius ^51,52^. These shared similarities are the reason that these two elements can both be incorporated into the pterin cofactor in some enzymes, such as *Rhodobacter capsulatus* DMSO reductase and nitrogenase ^6,53^. When producing *Pf*AOR recombinantly in a heterologous host that is not specific for W-enzyme production, this might lead to competition for the incorporation of W and Mo into the bis-MPT cofactor. To see if this is indeed happening in the recombinant AORs, the amount of Mo in the proteins was also measured.

When adding more tungstate to the medium, the percentage of incorporated molybdenum decreased from 29% in the 100 µM Na_2_WO_4_ sample to 10% in the 10 mM Na_2_WO_4_ sample. The higher tungstate concentration in the medium may have led to stronger competition with molybdenum for incorporation into the cofactor. Those results, together with the finding that a lower amount of sodium tungstate resulted in a higher incorporation of Fe (Fig. 2b), left us to conclude that we needed to use a higher amount of tungstate to prevent Mo incorporation. For that reason, all following protein expressions were performed using 1 mM Na_2_WO_4_ in the growth medium.

Finally, the supplementation of iron to the medium increased the iron incorporation into the protein, while the use of a higher IPTG concentration resulted in a lower incorporation of all metals involved (Fig. 2b).

The overall maximal W incorporation was 78% and Fe was 44%, which were obtained in two different samples. The native *Pf*AOR and GAPOR isolated from *P. furiosus* were used as positive control and contained 120 and 74% W and 117 and 106% Fe, respectively. Considering a standard deviation of approximately 15% for the determination of the protein concentration and 5-10% for metal concentration, the actual values will be closer to 100% for both proteins. Looking at the difference between native and recombinant proteins, the low level of activity using recombinant *Pf*AOR can, at least in part, be attributed to the substoichiometric incorporation of W and Fe.

### Expression and purification of ASR predicted proteins *Thermo*AOR3, *Thermo*AOR4 and *Thermo*AOR5

The expression of wild-type *Pf*AOR in *E. coli* Rosetta-Gami 2(DE3) resulted in a protein with low but measurable activity, along with substoichiometric pterin and metal incorporation. Changing the expression conditions of *Pf*AOR increased the incorporation of Fe and W but did not proportionally increase the activity. In an attempt to improve the heterologous expression of W-AORs to increase or maybe even alter the protein activity, we performed an ancestral sequence reconstruction (ASR) starting from *P. furiosus* AOR. ASR was performed using a phylogenetic tree comprising of 44 protein homologues and in total 5 evolutionary ancestors of AOR were predicted indicated by the labeled nodes. Node 1 resembles the node closest to *Pf*AOR, while Node 5 represents the ancestor furthest in the phylogenetic tree (Fig. S7). The extant sequences used for the prediction of Nodes 1 – 5 are all, but one, originating from archaea. The bacterial sequence is from *Thermovirga liennii* and may represent an example of lateral gene transfer given its high amino acid sequence identity to *Pf*AOR (73%). The other sequences predominantly represent archaea from the Phylum Methanobacteriota (homotypic synonym Euryarchaeota) and the class Thermococci. For the reconstruction of node 1 - 5 several archaea from the Phylum Thermoproteota (homotypic synonym Crenarchaeaota) and the class Thermoprotei are included. We therefore designate the ASR AORs as *Thermo*AOR numbered starting from *Pf*AOR, e.g. Node 1 being *Thermo*AOR1.

The predicted *Thermo*AORs share a sequence identity to *Pf*AOR ranging from 78 to 95%. The mutations present in the ASR proteins are predominantly located at the surface, for example close to the ferredoxin binding site (Fig. 3). The effect of some of these mutations can be seen by a small change in surface potential obtained using APBS (Adaptive Poisson-Boltzmann Solver), represented by red (negative) or blue colored surface (positive, Fig. 3a).

**Figure 3.**
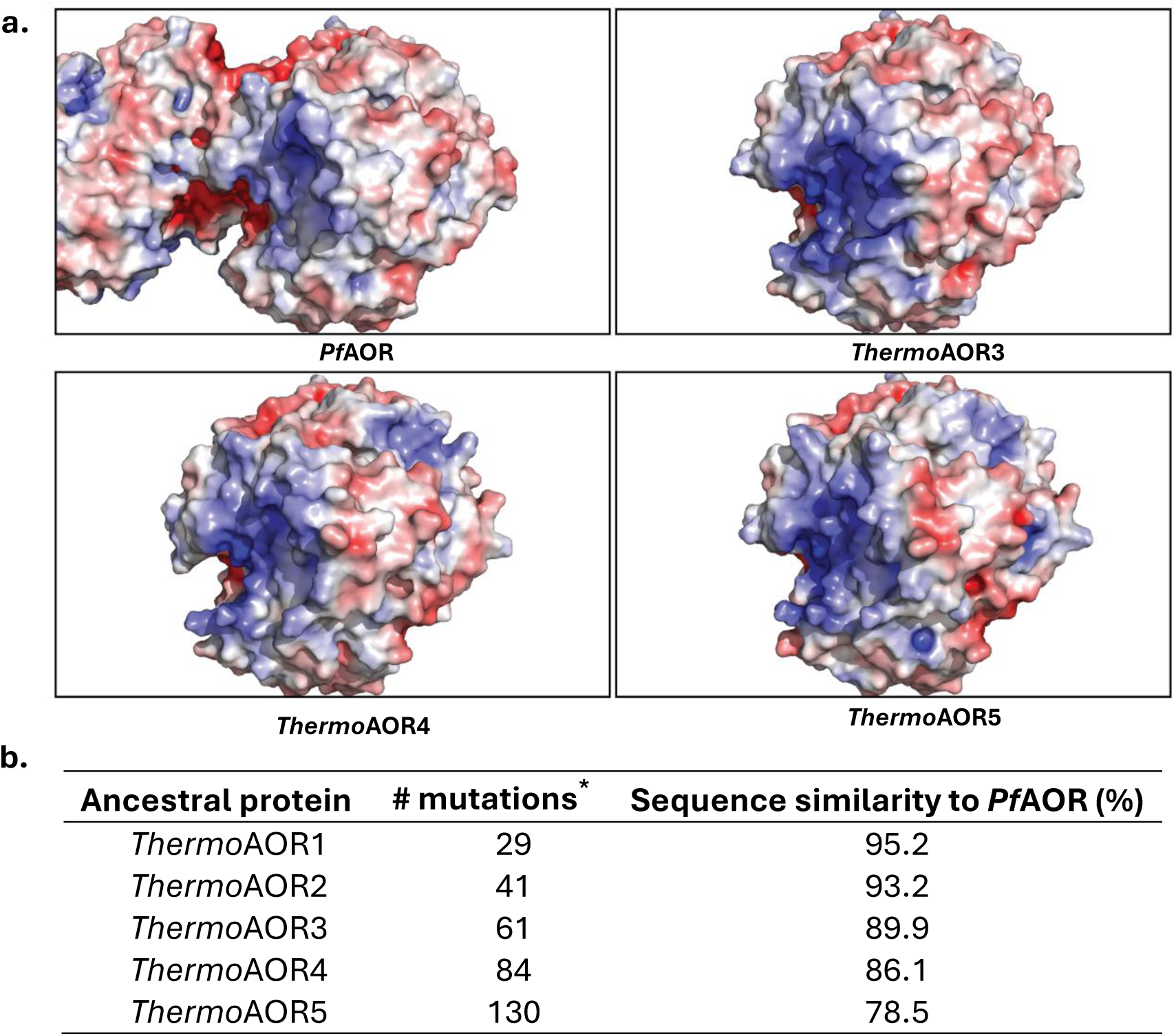
Sequence and charge similarity of predicted ancestral proteins compared to AOR. **a.** Surface charge around predicted ferredoxin binding site. Surface of the proteins is shown by charge distribution from negative (red) to positive (blue). **b.** Sequence similarity of ancestral proteins to *Pf*AOR. \**Pf*AOR has 605 amino acids.

AOR is believed to use a catalytic mechanism involving residues in a second coordination sphere of its metal cofactor. These residues are conserved among protein homologues and include Thr243, Glu313, Tyr427, and His448 (*Pf*AOR numbering). They play a role in catalysis as described by Szaleniec C Heider (2025, Fig. 4a) ^54^. In the ancestral variants, these proposed catalytic residues are conserved, as well as the distances between the pterin cofactor, [4Fe-4S] cluster and Arg76, a conserved residue found in all AOR-homologues (Fig. 4b) ^54^. The distances separating the different cofactors are similar to *Pf*AOR, allowing for efficient electron transfer ^55^. Due to the various mutations close to the protein surface, the surface charge near the substrate entry channel has been altered compared to the native *Pf*AOR protein, making the area less positively charged (Fig. 4c).

**Figure 4.**
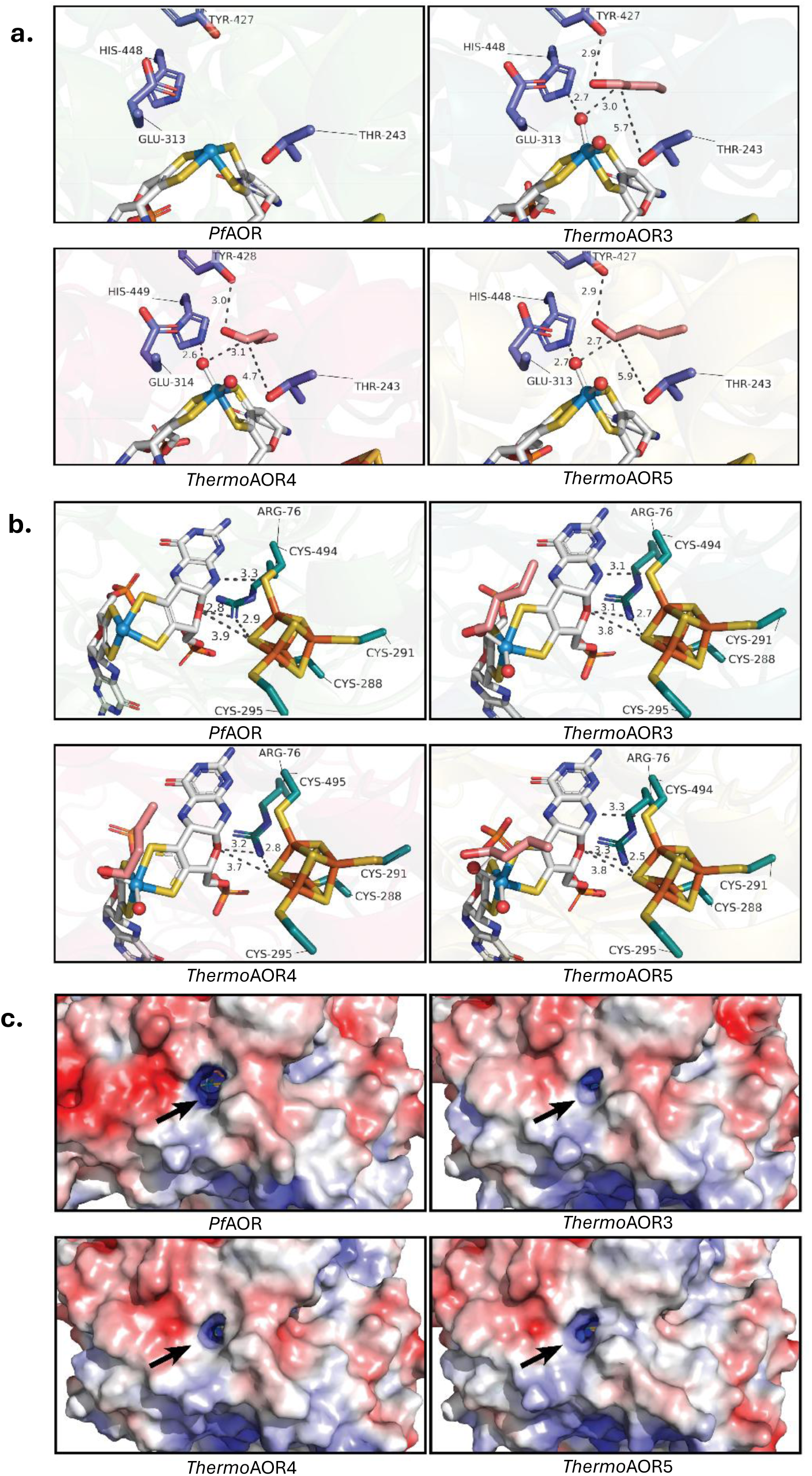
Active site structure of *Pf*AOR and predicted structures of *Thermo*AOR3, *Thermo*AOR4, and *Thermo*AOR5. **a.** Distances between the pterin cofactor, [4Fe-4S]-cluster and ARG76. **b.** Proposed catalytic residues in the active site. **c.** Surface charge around the entry channel of the proteins with negatively charged residues colored red and positive residues colored blue. Distances are represented by dashed black lines and given in Å.

To start the analysis of the ancestral proteins, *Thermo*AOR3, 4 and 5 were expressed in *E. coli* Origami 2(DE3) and purified using affinity chromatography. Successful soluble expression of *Thermo*AORs was confirmed using SDS-PAGE analysis, revealing prominent protein bands around 60-70 kDa, consistent with their expected molecular mass (Fig. S8). After purification, a yield of approximately 7 mg/L of each *Thermo*AORs was obtained.

### Ancestral sequence reconstruction resulted in an AOR variant with a 50-fold higher activity

The enzyme variants predicted with ASR represent evolutionary ancestors of *Pf*AOR ^56^. We therefore expect them to have the same, or similar biocatalytic properties as *Pf*AOR; the reversible oxidation of aldehydes into carboxylic acids, as well as high thermostability. As for *Pf*AOR, the enzyme activity of the ancestral enzymes was quantified using a spectrophotometric assay measuring oxidation of crotonaldehyde with benzyl viologen (BV) as artificial electron acceptor (Fig. 5c).

**Figure 5.**
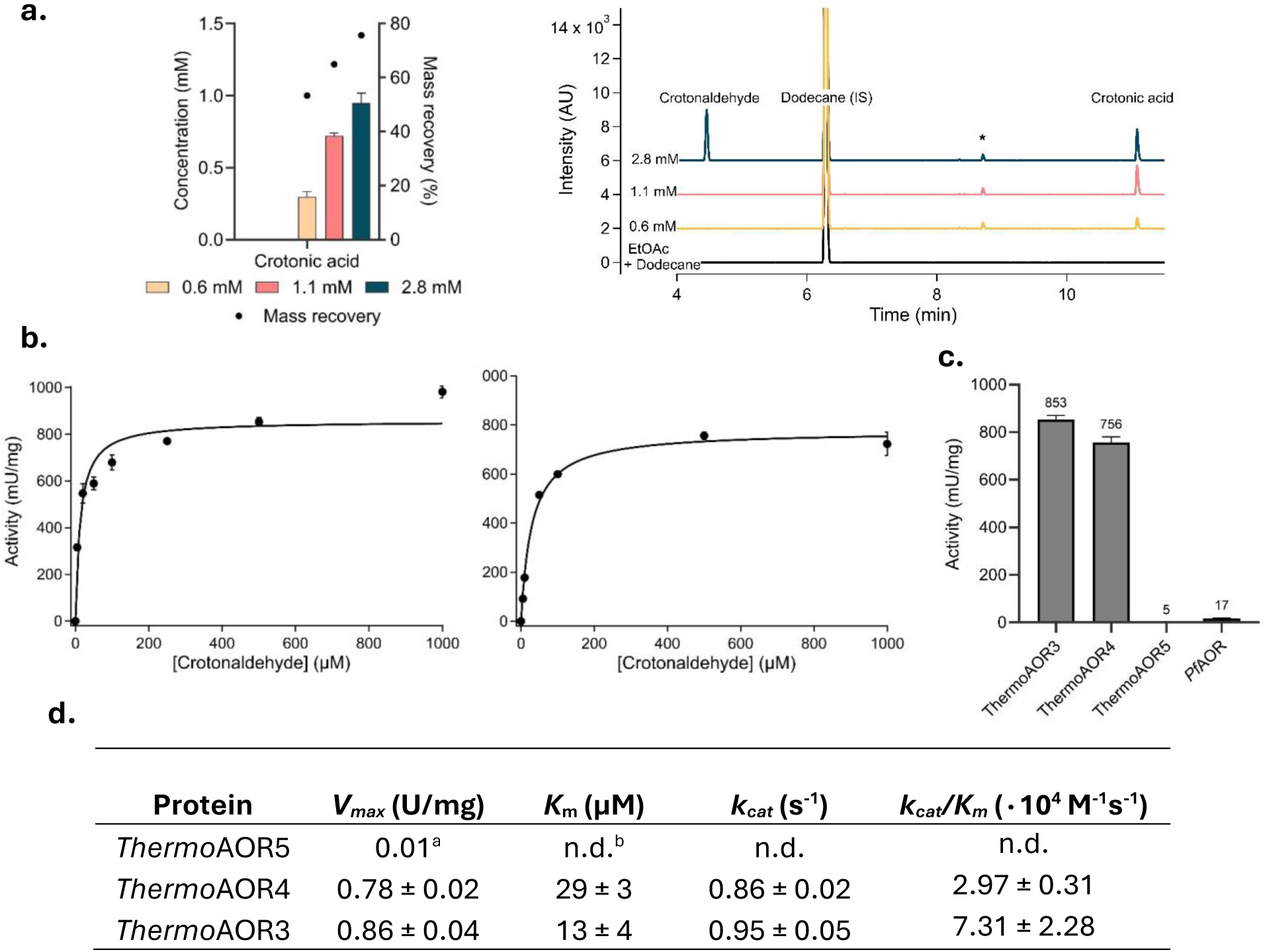
Activities of *Thermo*AOR3, *Thermo*AOR4 and *Thermo*AOR5. **a**. Left, GC quantification of *Thermo*AOR3 catalyzed oxidation of crotonaldehyde. The activity of *Thermo*AOR3 was tested using crotonaldehyde concentrations ranging from 0.6 to 2.8 mM at 60 °C in an anoxic environment for 4 hours. The reaction mixtures contained 3 mM BV, 0.6 – 2.8 mM crotonaldehyde and 0.42 nmol protein monomer with a total volume of 300 µL. Right, Gas chromatograms of crotonaldehyde oxidation by *Thermo*AOR3. Retention times; Crotonaldehyde: 4.46. n-Dodecane (IS): 6.30. DMSO: 10.15. Crotonic acid: 11.08. **b**. Michaelis-Menten kinetics of *Thermo*AOR3 (Left) and *Thermo*AOR4 (Right). Reactions contained 16 (*Thermo*AOR3) or 21 µg (*Thermo*AOR4) enzyme. **c**. Comparison of the activities of ASR proteins and recombinant wild-type *Pf*AOR. The reactions contained 12.7-21.2 µg protein. **d**. Kinetic properties of ancestral enzymes *Thermo*AOR3, *Thermo*AOR4 and *Thermo*AOR5. Values for *V_max_* and *K_m_* have been derived from a Michaelis-Menten fit of activity screening using crotonaldehyde and BV at 60°C. *k_cat_* is calculated per monomer using the *V_max_* values; ^a^ The initial rate at 1 mM crotonaldehyde is shown instead of the *V_max_*; ^b^ n.d., not determined. In **a**,**b**,**c**,**d** enzyme activity was measured at 60 °C in an anoxic environment as crotonaldehyde-dependent reduction of BV (ɛ_600_ = 7,400 M^-1^cm^-1^). The reaction mixture contained 3 mM BV and varying crotonaldehyde and enzyme concentrations in EPPS buffer (50 mM, pH 8.4 at 60°C). One unit (U) corresponds to 1 µmol of crotonaldehyde oxidized per minute.

The activity analysis of the ancestral enzymes showed that both *Thermo*AOR3 and *Thermo*AOR4 had a significant increase in activity compared to the previously expressed recombinant wild-type *Pf*AOR (Fig. 5c). *Thermo*AOR3 showed an oxidation activity of 853 ± 19 mU/mg, which is a 50-fold increase compared to recombinant wild-type *Pf*AOR. *Thermo*AOR4 showed a lower activity of 756 ± 24 mU/mg, representing a 44-fold increase compared to wild-type *Pf*AOR. *Thermo*AOR5, in contrast, showed a negligible crotonaldehyde oxidation activity of 5 mU/mg.

Of these three ASR proteins tested, *Thermo*AOR3 contains the least number of mutations compared to wild-type *Pf*AOR (61 mutations), followed by *Thermo*AOR4 with 84 mutations. *Thermo*AOR5 represents the evolutionary node furthest from the current *Pf*AOR protein with 130 mutations and a sequence identity of 78.5%. It is possible that this large number of mutations resulted in adverse structural changes, specifically close to the active site. Analysis of the predicted structure of *Thermo*AOR5, revealed two mutations in the active site close to the pterin cofactor that are not present in the other characterized *Thermo*AORs. The amino acids Ala92 and Lys450 in *Pf*AOR are mutated to Ser92 and Arg450 (Fig. S9). Being very close to the active site and cofactors, these mutations may negatively affect the protein activity.

The spectrophotometric activity assays measure the initial rate of aldehyde oxidation indirectly by measuring the rate of reduction of the electron acceptor BV. To verify the conversion of crotonaldehyde to crotonic acid by *Thermo*AOR3, gas chromatography (GC) analysis was performed to quantify the concentration of crotonaldehyde and crotonic acid (Fig. 5a).

GC analysis (Fig. 5a, Fig. S10) confirmed that *Thermo*AOR3 indeed converts crotonaldehyde to crotonic acid in the presence of the electron acceptor BV. Increasing the substrate concentration leads to an increased production of product, with full conversion obtained for the lower concentrations of crotonaldehyde used (0.6 and 1.1 mM) within 4h. The highest concentration of substrate (2.8 mM) was only partially oxidized to crotonic acid within 4h (Fig. 5a) because of rate limitation. Within 4h, 0.42 nmol of enzyme (monomer) produced approximately 0.3 µmol of crotonic acid, indicating a total number of turnovers of approximately 714. Since the catalytic activity of the enzymes is dependent on the W-content, expressing the turnovers related to the amount of incorporated W is a more reliable indication of the activity of intact protein (see ‘Cofactor composition of the ancestral *Thermo*AORs’ below). The total number of turnovers in 4h per W-cofactor for *Thermo*AOR3 under the conditions used here was approximately 5,000.

Within 4h incubation at 60°C approximately 34% of the crotonaldehyde was lost due to evaporation, which can be observed by the difference in crotonaldehyde present in the control samples before and after incubation (Fig. S10) and explains why the mass balance is incomplete (Fig. 5a).

### Michaelis-Menten kinetics of *Thermo*AOR3 and *Thermo*AOR4

As shown above, *Thermo*AOR3 and *Thermo*AOR4 showed a significant activity using 0.5 mM of crotonaldehyde of 853 and 756 mU/mg, respectively. To further characterize the catalytic properties, steady-state kinetics of crotonaldehyde conversion by *Thermo*AOR3 and *Thermo*AOR4 was measured (Fig. 5b). Analysis of the Michaelis-Menten curves showed an apparent *V_max_* and *K*_m_ for *Thermo*AOR3 with crotonaldehyde and 3 mM BV of 857 ± 36 mU/mg and 13 ± 3.8 µM, respectively. *Thermo*AOR4 shows an apparent *V_max_* of 776 ± 19 mU/mg and *K*_m_ of 29 ± 3.3 µM at 60°C (Fig. 5b).

These results show that both enzymes have a similar substrate dependence of the activity characterized by the comparable *K*_m_ values. When looking at the *V_max_* values, *Thermo*AOR3 shows a higher *V_max_* than *Thermo*AOR4 (857 to 776 mU/mg, respectively). It is worth noting that the activity of *Thermo*AOR4 gradually decreased upon repeated freeze-thaw cycles and therefore all activity measurements were normalized to the initially observed activity of 756 mU/mg using 0.5 mM crotonaldehyde. The gradual reduction in activity is most likely the result of sensitivity to multiple freeze-thaw cycles.

The *K_m_* of *Thermo*AOR4 and *Thermo*AOR3 at 29 and 13 µM respectively, are close to the reported *K*_m_ at 40 µM of native *Pf*AOR for crotonaldehyde (Fig. 5d). The reported *V_max_* of native *Pf*AOR for crotonaldehyde is 67 U/mg at 80°C using methyl viologen (MV) as electron acceptor ^12^. Assuming a two-fold lower rate per 10°C, the *Pf*AOR *V_max_* is approximately 17 U/mg at 60°C ^12^. The *V_max_* of native *Pf*AOR is substantially higher than the *V_max_* of the ancestral enzymes, consistent with their substoichiometric W-cofactor and [4Fe-4S] cluster incorporation (Table 2). To assess whether the temperature optimum of *Thermo*AOR3 is similar to native *Pf*AOR, enzyme activity using crotonaldehyde was measured at different temperatures ranging from 40 to 90°C (Fig. 6).

**Figure 6.**
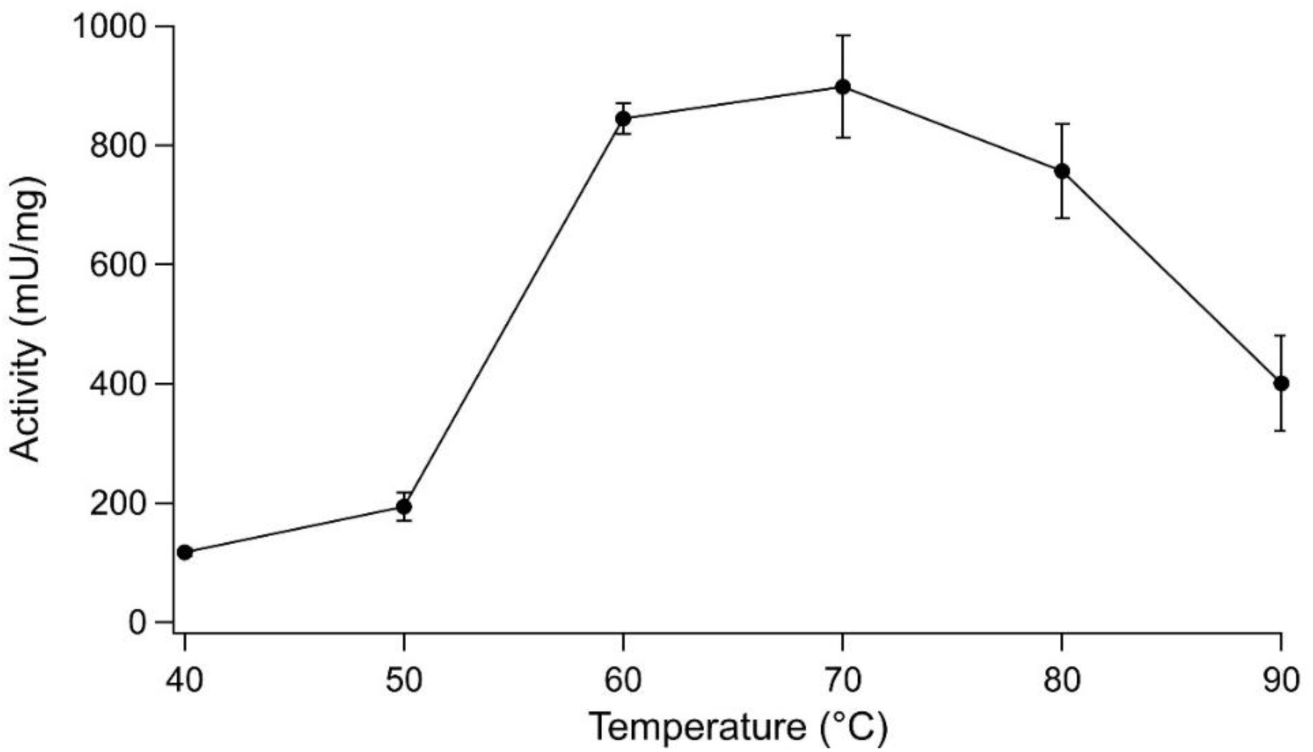
Temperature profile of *Thermo*AOR3 oxidation activity using crotonaldehyde. Enzyme activity was measured at 40-90°C in an anoxic environment using benzyl viologen as artificial electron acceptor. The reactions contained 3 mM BV, 0.5 mM crotonaldehyde and 14 µg enzyme in EPPS buffer (50 mM, pH 8.4 at 60°C). One unit (U) corresponds to 1 µmol of crotonaldehyde oxidized per minute.

**Table 2.** Cofactor content of purified *Thermo*AOR3, *Thermo*AOR4 and *Thermo*AOR5.

| Protein | Fe<sup>a</sup> (mol/mol (%)<sup>c</sup>) | Mo<sup>a</sup> (mol/mol) | W<sup>a</sup> (mol/mol (%)<sup>c</sup>) | Pterin<sup>b</sup> (mol/mol (%)<sup>c</sup>) |
|---|---|---|---|---|
| *Thermo*AOR5 | 0.06 (1.5%) | 0.01 | 0.10 (10%) | 0.18 (9%) |
| *Thermo*AOR4 | 0.54 (14%) | 0.02 | 0.07 (7%) | 0.14 (7%) |
| *Thermo*AOR3 | 0.69 (17%) | 0.02 | 0.15 (15%) | 0.17 (9%) |
| BSA | 0.01 (n.a.<sup>d</sup>) | 0.00 | 0.00 | n.d.<sup>e</sup> |
| Xanthine oxidase | n.d. | n.d. | n.d. | 1.00 (100%) |

<sup>a</sup> Metal content has been measured using an ICP-MS analysis and bovine serum albumin (BSA) was included as negative control; <sup>b</sup> Pterin content was measured with a fluorometric assay with an excitation wavelength of 350 nm and emission wavelength of 450 nm. Xanthine oxidase was included as positive control for the pterin determination and all data was normalized to a xanthine oxidase content of 1 mol pterin/monomer; <sup>c</sup> Values are also shown as percentage of maximally expected cofactor content of 4 mol/mol Fe, 1 mol/mol W and 2 mol/mol pterin. The average error of the ICP-MS measurements was; Fe – 0.04 mol/mol, Mo – 0.001 mol/mol, W – 0.002 mol/mol. The pterin measurements had an error of 0.04 mol/mol on average; <sup>d</sup> n.a., not applicable; <sup>e</sup> n.d., not determined.

The activity of *Thermo*AOR3 increases from 40°C, with an optimum at circa 70°C (Fig. 6). At temperatures above 70°C, the activity decreases to approximately 750 mU/mg and further to 400 mU/mg at 90°C. Native *Pf*AOR has been reported to continuously increase activity from 25 to 100°C, with a plateau at 98°C of approximately 100 U/mg ^16^. This shows that the ancestral variant of *PfA*OR, *Thermo*AOR3, is not as thermostable as the extant enzyme and therefore has a lower apparent temperature optimum.

Interestingly, we discovered that the lysis method heavily influences the maximal activity obtained and the temperature stability of the protein (Fig. S11). When changing the cell lysis method from a chemical lysis using FastBreak™ lysis cell reagent to lysis using sonication, the activity obtained increased from 0.86 U/mg to 2 U/mg at 60°C. When measuring the activity of *Thermo*AOR3 lysed using sonication at increasing temperatures from 40 to 90°C, a continues increase in initial rate was observed, similar to native *Pf*AOR (Fig. S11). This indicates that the (chemical) lysis of the protein might influence the overall activity and thermostability of the protein and improvements in activity upon optimization of the culture and purification conditions remain possible.

### Cofactor composition of the ancestral *Thermo*AORs

AORs use a single W-bis-MPT cofactor and single [4Fe-4S]-cluster per subunit to catalyze the two-electron oxidation of aldehydes. In each protein monomer, 1 mol W, 4 mol Fe and 2 pterin moieties are expected to be present if all functional groups are properly incorporated. To test this, the metal content of the purified proteins was analyzed using ICP-MS analysis and the pterin content was determined using a fluorescence assay measuring the fluorescent derivative of the bis-MPT cofactor, with pterin-6-carboxylic acid as standard curve.

For the ICP-MS analysis, bovine serum albumin was included as a negative control and was found to contain no detectable amounts of W and Mo and an iron content of 0.01 mol/mol (Table 2). Of all ASR enzymes, *Thermo*AOR3 contained the highest amount of Fe (0.69 mol/mol) followed by *Thermo*AOR4 (0.54 mol/mol) and *Thermo*AOR5 (0.06 mol/mol). All values are significantly lower than the theoretically expected 4 mol Fe per monomer, indicating incomplete [4Fe-4S]-cluster incorporation. The low iron content of *Thermo*AOR5 compared to the other two ancestral proteins could also play a part in the negligible activity obtained using this enzyme. The catalytically most active ASR enzyme, *Thermo*AOR3, contained the highest amount of W, 0.15 mol/mol, approximately 15% of the expected 1 mol W. In all proteins, negligible amounts of Mo were found of maximally 0.02 mol/mol, indicating a successful exclusion of Mo from the cofactor. The pterin content is comparable in all ASR proteins, with 7-9% of the expected amount incorporated in all proteins. This incorporation is also in accordance with the amount of incorporated W (approximately 7-15% for all).

Taking the iron, W and pterin content into account, approximately 10-20% of both iron-sulfur cluster and W-cofactors have been incorporated in *Thermo*AOR3. Given that only intact protein (approx. 15%) can catalyze the oxidation of crotonaldehyde, functional *Thermo*AOR3 showed approximately 5,000 turnovers per 4h per W-cofactor based on the aforementioned GC analysis results (Fig. 5a) and we can estimate a 6.7-fold higher *V_max_* of 5.7 U/mg for the holoenzyme *Thermo*AOR3.

It is notable that the analysis of the pterin and metal content showed that the ASR did not produce a protein with a higher W-cofactor and [4Fe-4S] cluster content than the recombinantly expressed wildtype *Pf*AOR] (Fig. 2b). ASR leads to significantly more active enzymes, with an activity increase of approximately 50-fold. This increase in activity therefore has to be caused by other factors than cofactor incorporation, which we are currently examining.

Since the Fe incorporation was incomplete, we further investigated whether *Thermo*AOR3 contained complete [4Fe-4S] clusters using EPR (electron paramagnetic resonance) and UV-vis analysis (Fig. 7).

**Figure 7.**
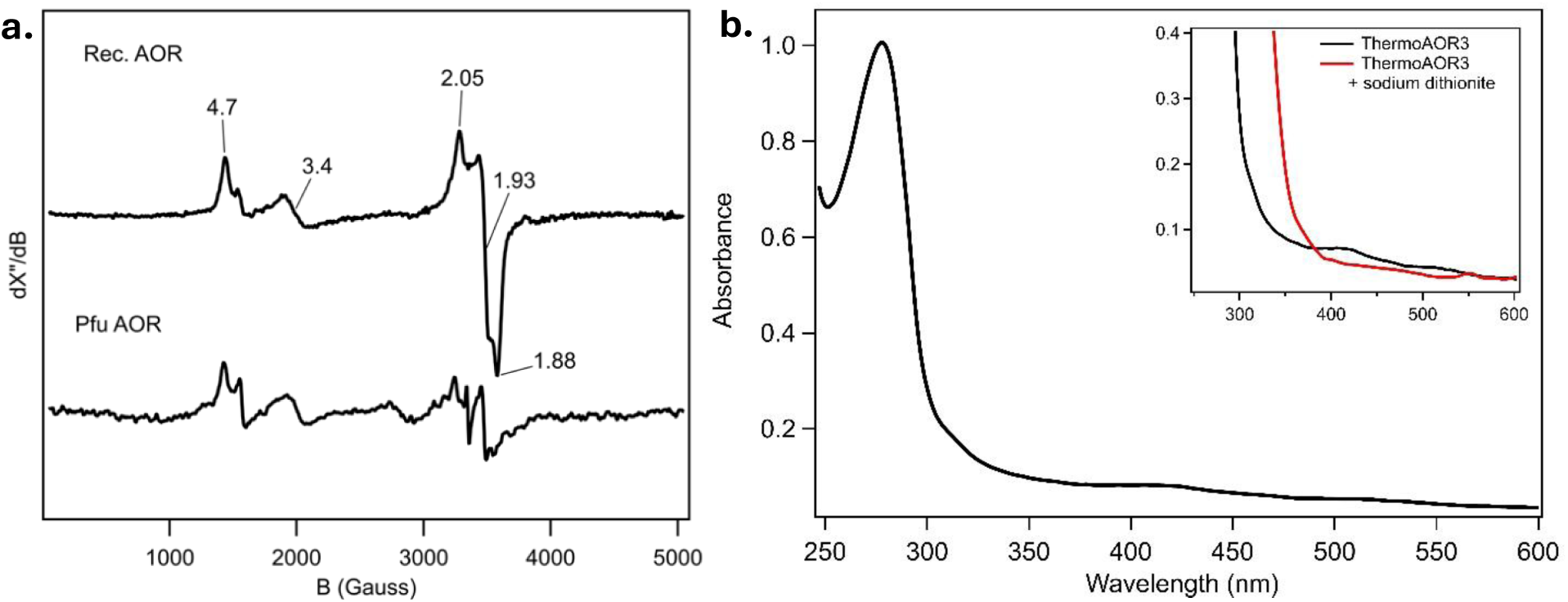
EPR and UV-Vis analysis of *Thermo*AOR3. **a**. Comparison of EPR spectra of as-isolated native *Pf*AOR and as-isolated recombinantly produced *Thermo*AOR3 (Rec. AOR). Spectra acquisition was performed under microwave frequency of 9.403 GHz, microwave power 20 mW, a modulation frequency of 100 kHz, and a modulation amplitude of 10 G. T=15 K. **b**. UV-Vis analysis of as-isolated *Thermo*AOR3. **Inset**. Comparison of UV-Vis analysis of *Thermo*AOR3 and *Thermo*AOR3 reduced with 250 µM sodium dithionite.

Spectroscopic analysis of *Thermo*AOR3 showed a broad absorbance band between 300-500 nm, which could be reduced upon the addition of sodium dithionite. This band is characteristic for iron-sulfur clusters in proteins and therefore indicates the presence of a redox active FeS cluster ^57^. Moreover, EPR spectroscopy of the as-isolated enzyme showed the S=1/2 and S=3/2 signals with identical g-values to those of the [4Fe-4S]^+^ cluster of native *Pf*AOR (Fig. 7a) ^58^.

### Ancestral *Thermo*AOR3 has a broad substrate scope for aldehyde oxidation

*Pf*AOR is a member of the AOR-family of enzymes that are all able to convert either one specific or various different aldehydes. Members of this family have been found to contain different substrate specificities. GAPOR, for example, only catalyzes the conversion of its physiological substrate D-glyceraldehyde-3-phosphate (D-GAP), whereas other members, such as *Pf*AOR exhibit a much broader substrate range ^16^. Interestingly, *Pf*AOR cannot catalyze the oxidation of D-GAP and has very low activity towards hexanal, the preferred substrate of WOR5, another W-AOR family enzyme from *P. furiosus* ^15^.

To test if *Thermo*AOR3 has a similarly broad substrate scope as *Pf*AOR and analyze if it is also capable of converting substrates preferred by other AOR-family enzymes, the activity of *Thermo*AOR3 for a range of aldehydes was screened. These substrates include the aforementioned D-GAP and hexanal, preferred by the AOR-family enzymes GAPOR and WOR5, respectively ^15^. The other tested substrates range from linear aliphatic (e.g. butyraldehyde), branched chain aliphatic (e.g. isobutyraldehyde), aromatic (e.g. vanillin) to heterocyclic (furfural) aldehydes. Moreover, the capability of *Thermo*AOR3 to catalyze the thermodynamically challenging carboxylic acid reduction was measured using acetate as substate. The activities of all substrates are compared to the activity obtained using the reference substrate crotonaldehyde and are therefore normalized to this activity (Fig. 8a).

**Figure 8.**
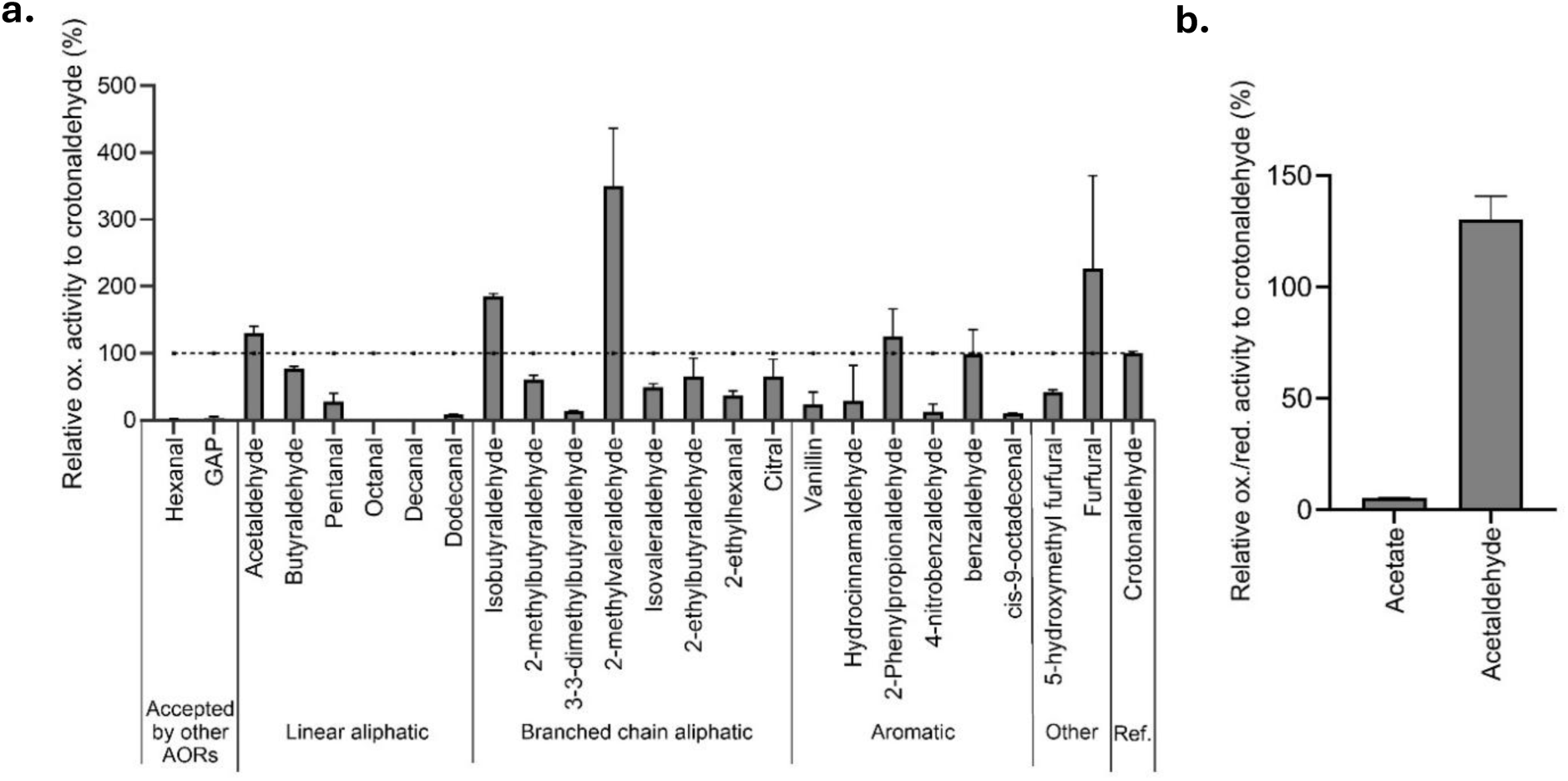
Substrate specificity of *Thermo*AOR3. Activity of *Thermo*AOR3 was measured for different aldehydes (**a**) and one carboxylic acid, acetate (**b**). Oxidation reactions contained 3 mM BV, 0.5 mM substrate and 9.5 µg enzyme in 50 mM EPPS buffer pH 8.4. Activity was measured as the initial rate at 60°C in an anoxic environment by following the reduction of BV (ɛ_600_ = 7.4 mM^-1^cm^-1^). The carboxylic acid reduction reaction contained 5.1 mM HMV, 4.6 mM Ti(III)citrate, 10 mM phenylhydrazine, 60 mM acetate, and 10.8 µg enzyme in 100 mM KPi buffer pH 6.0. Crotonaldehyde served as a reference substrate and was set at 100% activity. The activities of the other substrates are normalized to the activity with crotonaldehyde.

*Thermo*AOR3 showed little to no activity when using the substrates accepted by other AOR family enzymes GAPOR and WOR5, D-GAP and hexanal, just as for native *Pf*AOR. The short chain aliphatic aldehydes butyraldehyde and acetaldehyde were oxidized with an activity comparable to crotonaldehyde (set at 100%) or higher (78 and 130%, respectively). Increasing the chain-length to pentanal and further to octanal and decanal reduced the activity significantly, with octanal and decanal having no measurable activity. An even longer chain length (C12, dodecanal) showed a low, but measurable relative activity of 8%. Generally, *Thermo*AOR3 showed a higher specific activity towards branched-chain aliphatic aldehydes. Adding a methyl group at the C2 position of pentanal (C5), forming 2-methylvaleraldehyde (C6), increased the activity significantly from 28 to 350%, resulting in the highest activity measured for all compounds tested. In industry, 2-methylvaleraldehyde is used as a precursor for the production of several pharmaceuticals including the sedative meprobamate, indicating its relevance to be used in future industrial applications ^59^. Surprisingly, for the branched chain aldehydes, longer chain compounds (>C5) were still accepted and oxidized with a relatively high activity, e.g. 37% for 2-ethylhexanal (C8) and 66% for citral (C10). Even an unsaturated fatty aldehyde, cis-9-octadecenal (C18), used as a model compound to explore the acceptance of long chain fatty acids, was converted with a low, but measurable activity of 10%. Increasing the size of the branch was tested using 2-methylbutyraldehyde (61%) and 2-ethylbutyraldehyde (66%). The similar rate suggests a bulkier side chain (from C1 to C2) did not hinder the interaction of the compounds with the active site. Adding two side chains using 3-3-dimethylbutyraldehyde, decreased the activity compared to 2-methylbutyraldehyde containing only one methyl side chain from 61 to 13%.

Aromatic aldehydes were converted with varying efficiencies, ranging from 126% for 2-phenylpropionaldehyde to only 13% when using 4-nitrobenzaldehyde. The heterocyclic aldehyde furfural and 5-hydroxymethylfurfural (HMF), industrially relevant building blocks for the production of specialty chemicals, were also accepted. Using furfural resulted in a relatively high activity of 227%, while the use of HMF showed a lower activity of 41% ^60^.

### *Thermo*AOR3 catalyzes carboxylic acid reduction

Finally, the capability of *Thermo*AOR3 to catalyze the thermodynamically challenging reverse reaction, the reduction of carboxylic acids was determined. The in-vitro carboxylic acid reduction activity converting acetate was 40 ± 3 mU/mg at 60°C using reduced HMV (*E^’0^* = −610 mV vs. SHE) as electron donor and phenylhydrazine as an aldehyde trapping agent (Fig. 8b, Fig. S12) ^35^. This is 3.6% of the activity obtained when screening *Thermo*AOR3 in the oxidative direction using acetaldehyde. Previous reports show that other native AORs exhibit a significantly slower reduction than oxidation reaction, consistent with the unfavorable equilibrium. During the screening of the carboxylic acid reduction activity of *Clostridium thermoaceticum* AOR, the oxidation activity was found to be 20 times faster than the reduction activity using methyl viologen (MV) as electron donor at pH 5.5 with 20 mM semicarbazide to trap the produced aldehyde and 60 mM propionate as substrate. For *Thermococcus paralvinellae* (previously named *Thermococcus* ES-1) AOR, the highest acetate reduction activity was shown to be approximately 5% of the aldehyde oxidation activity as well using MV as electron donor at pH 5.5 and using 60 mM acetate as substrate ^11,34^.

### Conclusions

In this study we produced a functionally active variant of AOR by designing and expressing ancestral nodes of *Pf*AOR predicted using ancestral sequence reconstruction. Three of these ancestral nodes have been characterized and two of these proteins, *Thermo*AOR3 and *Thermo*AOR4 have an approximately 50-and 44-fold higher activity compared to recombinantly expressed wild-type *PfAOR* in *E. coli* using optimized strains and cultivation conditions. To our knowledge, this study demonstrates the first recombinant expression of a functional W-AOR, representing an exclusively W-containing enzyme family. The ancestral enzymes have substoichiometric incorporation of the W-cofactor, as well as the [4Fe-4S] cluster needed for enzyme catalysis. This indicates the expression and incorporation of the cofactors can still be improved by for example co-expressing chaperones or specific W-cofactor biosynthesis proteins, thereby increasing enzymatic activity. To achieve this, a strategy could be engineering of the *E. coli* strain to introduce specific W-cofactor and FeS-cluster biosynthesis machinery or using a co-expression system as described by Phan et al. (2026) for the production of W-FDH in *E. coli ^c1^*. Using these systems, the production and incorporation of the cofactors could be improved, which ultimately could lead to improved catalytic activity. The fact that even with substoichiometric incorporation of the cofactors, the enzymatic activity was significantly increased compared to the recombinant wild-type enzyme, suggests that the cofactor biosynthesis capability of *E. coli* was not the only bottleneck in the production of functional W-enzymes.

Using ASR, amino acid substitutions have been implemented by a statistical analysis of each amino acid position in the protein sequence considering the variety of the extant sequences in the phylogenetic tree. Despite the fact that no restraints are given to conserve either the active site or entry channels, we observe that the mutations are introduced all over the protein leaving the active site and channels intact^62^.

*Thermo*AOR3 is, just as *Pf*AOR, able to convert a variety of aldehydes, with the highest activities observed using branched chain aliphatic aldehydes and short chain linear aliphatic aldehydes. Using crotonaldehyde as model substrate, *Thermo*AOR3 and *Thermo*AOR4 showed Michaelis-Menten kinetics with a *V*_max_ approximately 5% of the value reported for native *Pf*AOR and a *K*_m_ comparable to the *Pf*AOR *K*_m_. Next to that, the enzyme is also able to catalyze the thermodynamically challenging reduction of acetate to acetaldehyde using reduced hexamethylviologen as electron donor.

Just as *Pf*AOR, *Thermo*AOR3 has a higher temperature optimum, but does not seem to be as thermostable as *Pf*AOR. The optimum temperature lies around 70°C to 90°C depending on the enzyme isolation method. The isolation method also affected the maximal specific activity that was obtained with 2 U/mg observed as the highest activity in any preparation. This suggests that further improvements of the activity upon improving purification conditions can still be made. Finally, *Thermo*AOR3 contained approximately 10-15% of the maximally expected pterin, W and Fe and was shown to have a redox active [4Fe-4S] cluster.

This work represents an important breakthrough that will make this unique class of W enzymes available to the field of biocatalysis and industrial biotechnology by enabling scalable enzyme production and application of the full extent of modern enzyme engineering tools. Furthermore, it will accelerate mechanistic investigations for these enzymes and allow mining of the largely unexplored biodiversity of the W-AOR family.

## Supporting information

Supplementary Chai-1 structure

Supplementary Chai-1 structure

Supplementary Chai-1 structure

## Acknowledgements

The authors acknowledge funding from the European Union’s Horizon Europe Innovation Council under grant agreement No 101129798, EIC PathFinder open project W-BioCat. PLH acknowledges support from COST Action FeSImmChemNet (CA21115) supported by COST (European Cooperation in Science and Technology) and funding from the European Union’s Horizon Europe research and innovation programme under grant agreement no. 101183014 (project McGEA). We also want to thank the Department of Radiation Science and Technology, Delft University of Technology, and specifically Özlem Chalashkan for setting up the analysis methods to measure W and performing the ICP-MS experiments in their facilities. We thank Alice Pedroni (Department of Biotechnology, Delft University of Technology) for performing the synthesis of HMV.

## Author contributions

D.B., Y.W., P.W., R.S. and P.-L.H. designed and performed the experiments and data evaluation. I.M. and J.V. performed the ancestral sequence reconstruction. P.L.H., F.H., I.M. and J.V. conceived the project. D.B. and P.-L.H. wrote the manuscript and all authors jointly edited the manuscript. All authors approved the final version of the manuscript.

