## Supplementary info for "Ancestral sequence reconstruction enables functional expression of archaeal thermostable tungsten-containing aldehyde:ferredoxin oxidoreductases in *E. coli*"

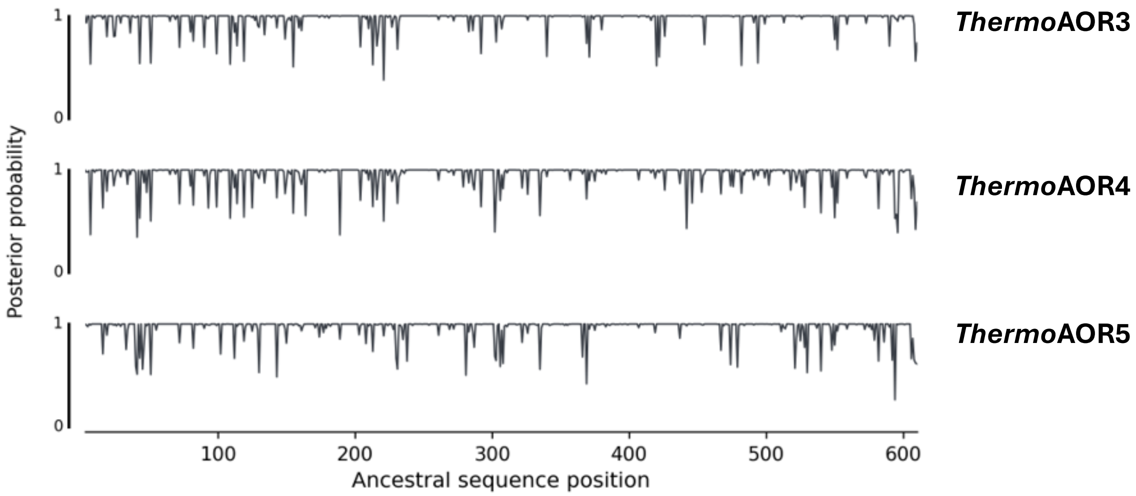

**Figure S1. Posterior probability profiles of ThermoAOR3, ThermoAOR4, and ThermoAOR5.**

Posterior probability profiles of the inferred amino acids across the 610 positions for the targeted internal nodes. The amino acid incorporated at each position of the ancestral protein is the one with the highest posterior probability. Mean posterior probabilities for *Thermo*AOR3, *Thermo*AOR4 and *Thermo*AO5 are 0.97, 0.96 and 0.96, respectively.

**Table S1. DNA sequences of gene constructs codon optimized for expression in *E. coli*.**

| Protein | DNA sequence |
|---|---|
| *Pf*AOR | ATGGTGTACGGAAACTGGGGGCGCTTCATCCGTGTGAATTTGTCCACTGGCGACATCAAAGTCG<br>AAGAGTACGACGAGGAACTTGCCAAGAAGTGGTTGGGAAGTCGTGGCCTGGCAATCTATCTGCT<br>GCTGAAAGAAATGGATCCCACCGTTGACCCCTTATCTCCTGAAAATAAGTTGATCATTGCAGCG<br>GGCCCGTTGACCGGGACCAGTGCTCCTACAGGAGGACGCTACAATGTAGTAACAAAGTCACCTC<br>TTACAGGTTTCATTACTATGGCAAACTCGGGTGGTTACTTCGGGGCAGAGTTAAAGTTTGCCGG<br>TTACGACGCCATTGTCGTAGAGGGAAAGGCGGAAAAGCCGGTTTATATTTACATTAAAGATGAA<br>CACATTGAAATTCGTGATGCCTCGCATATCTGGGGGAAAAAGGTAAGTGAAACCGAAGCTACGA<br>TCCGCAAAGAAGTCGGTTCAGAAAAGGTCAAAATTGCGTCAATTGGCCCCGCAGGGGAAAACCT<br>GGTGAAATTCGCTGCCATTATGAATGATGGGCATCGTGCTGCGGGGCGCGGCGGAGTAGGGGCT<br>GTTATGGGGTCGAAGAATTTAAAAGCCATCGCAGTAGAAGGGAGCAAAACAGTGCCGATCGCGG<br>ATAAACAAAAATTTATGTTAGTAGTCCGCGAAAAGGTTAATAAACTGCGTAATGACCCGGTCGC<br>GGGCGGGGGGTTACCGAAGTACGGCACGGCTGTATTGGTTAATATTATCAACGAGAATGGTTTG<br>TATCCTGTTAAAAACTTCCAGACGGGTGTCTACCCATACGCGTACGAACAATCCGGAGAGGCAA<br>TGGCAGCAAAGTATTTGGTACGTAATAAACCTTGCTATGCTTGTCCAATTGGGTGTGGGCGTGT<br>GAACCGCCTGCCTACTGTCGGTGAAACTGAAGGACCGGAATACGAGTCAGTCTGGGCGCTTGGG<br>GCCAACTTAGGAATTAACGACTTGGCTTCTATCATCGAGGCCAACCATATGTGTGATGAATTAG<br>GTCTGGACACGATCTCTACAGGGGGTACTTTGGCTACGGCAATGGAGCTTTACGAAAAGGGACA<br>CATTAAGGATGAAGAATTAGGGGACGCACCCCCATTTCGCTGGGGAAACACGGAAGTATTGCAC<br>TATTATATCGAGAAGATTGCAAAACGCGAGGGATTCGGAGACAAGCTGGCGGAAGGATCCTACC<br>GCTTGGCCGAGAGTTACGGCCACCCGGAGTTGAGCATGACGGTTAAAAAACTTGAGTTACCGGC<br>CTATGACCCCCGTGGTGCAGAAGGCCACGGTTTGGGGTATGCCACTAATAATCGCGGTGGGTGT<br>CACATTAAGAACTATATGATCTCACCAGAGATCTTAGGGTACCCTTATAAGATGGACCCACACG<br>ACGTGTCGGATGATAAAATCAAAATGTTAATTTTATTTCAGGACCTGACGGCGCTGATTGACTC<br>AGCAGGCTTGTGCTTGTTTACTACATTTGGACTGGGGGCTGACGATTACCGCGATTTATTAAAT<br>GCGGCCCTTGGTTGGGACTTCACTACCGAAGATTATTTGAAGATTGGGGAACGCATTTGGAATG<br>CCGAACGCTTGTTTAATCTTAAAGCGGGGCTTGACCCAGCCCGTGACGACACCTTGCCCAAACG<br>TTTTCTTGAGGAACCTATGCCGGAAGGCCCCAACAAGGGGCACACAGTACGTTTGAAGGAAATG<br>TTGCCTCGTTACTATAAATTACGCGGCTGGACGGAAGATGGCAAAATCCCAAAGGAGAAACTGG<br>AGGAGTTGGGAATCGCTGAGTTTTACTGAAGCTT |
| *Thermo*AOR5 | ATGATGGGCTATATGGGCAAAATTCTGCGCGTGAATCTGACCACCGGCGAAATCAAGGTCGAAG<br>AACTGGATGAAGAACTGGCGAAAAAATTTATTGGCGGCCGCGGTCTGGCCAGCAAATACCTGTA<br>TGATGAGATTGATCCGACCGTGGACCCGCTGTCGCCGGAAAACAAACTGATTTTTGCCACCGGC<br>CCGCTGACCGGCACCACCGCACCGACCGGCGGCCGCTACATGGTGGTAACCAAAAGCCCGCTGA<br>CCGGCTTCATTGCCAGCAGTAATTCAGGCGGTTTTTTTGGCCCGGAACTGAAATTTGCGGGCTA<br>CGATATGCTGATTGTGGAAGGCAAAGCGGATAAACCGGTATATATTAGCATTAAAGATGACAAC<br>GTGGAAATTAAAGATGCGAGCCATCTGTGGGGCAAAAAAGTGAGCGAAACCACCGATATTTTAC<br>TGGAAGAATTCGGCGATAAAAAAGCGCGTGTGGCCTGTATTGGCCCGGCCGGTGAAAACCTGGT<br>TAAATTTGCCGCGATTATGAATGACAAACACCGCGCGGCCGGCCGTTCTGGCGTTGGCGCCGTT<br>ATGGGCAGTAAAAACCTGAAAGCCATTGTGGTGCGTGGTAGCAAAAAAGTGGAAGTTGCCGATA<br>AAGAAAAATTTATGGAAGTTGTGAAAGAAAAAATTAAAAAGATTAAAGAAAATCCGGTGACGGG<br>AGGTGGCCTGCCGAAATACGGCACCGCGGTGCTGGTGAACATTATTAACGAAAACGGTCTGTAT<br>CCGACGCGCAACTTTCAAACCGGCGTATTTGAATATGCGTACGAGCAGAGCGGCGAAGCGCTGG<br>CGGCGAAATACCTGGTGCGTAACAAGCCGTGCTTTGCGTGCCCGATTGGCTGCGGCCGCGTGAG<br>CAAACTGCCGGCGGTGGGCGAAACCGAAGGCCCGGAGTATGAAAGCATTTGGGCGCTGGGCGCG<br>AACTGTGGCATTAATGATCTGGCGAGCATTGTGGAAGCGAATCATCTGTGCGATGAATACGGCC<br>TGGATACAATTTCAACGGGCGGCACCCTGGCAGCAGCGATGGAACTGTATGAAAAAGGCATCAT<br>TAAAGAAGAAGAACTGGGCGATGGCCCCCCGTTACGTTTTGGCAACAGCGAAGTGCTGCACTAT<br>TATATCGAGAAAATTGCCAAACGCGAAGGCTTCGGTGATAAACTGGCGGAAGGCAGTTATCGCC<br>TGGCCGAAAGCTATGGTCATCCGGAATATAGCATGAGCGTGAAAAAACAGGAATTACCGGCGTA<br>TGATCCGCGCGGCGCCGAAGGCCATGGCCTGGAATACGCGACCAGCAATCGTGGCGGCTGCCAT<br>GTGCGCGGCTATATGATTAGCCCGGAGATTCTGGGCGTGCCGAAAAAACTGGACCCGCATGACA<br>TTAGCGAAGAAAAAGCGAAATGGGTGAAAATTTTTCAGGATCTGACGGCGGTGATTGATAGCGC<br>AGGTCTGTGCCTGTTTACCAGCTTCGCCCTGGGCGCGGATGATTATCGTGATCTGCTGAATGCG |

| | |
|---|---|
| | GCGACCGGCTTTGATTATACCACCGAAGAAATGATGAAAGCGGGCGAACGCATTTGGAATCTGG<br>AACGCCTGTTCAATCTGAAAGCCGGTCTGGACCCGGAAAAAGATGACACCCTGCCGAAACGTTT<br>CCTGGAAGAACCCATGCCGGAAGGCCCGAATAAAGGCCATGTGGTGCGCCTGAAAGAACTGCTG<br>CCGAAATATTATAAACTGCGTGGCTGGAACGAAGATGGCATTCCGACCGAAGAAAAACTGAAAG<br>AACTGGGCCTGGCCGAATTTTACTAA |
| *Thermo*AOR4 | ATGATGGGCTACATGGGCAAAATTCTGCGTGTTAATCTGACGACCGGCGAAATTAAAGTGGAAG<br>AACTGGATGAAGAACTGGCGAAAAAATGGATTGGCGGTCGCGGCCTGGGCACCTATTATCTGCT<br>GAAAGAGATGGACCCCAAAGTTGATCCGCTGTCGCCGGAAAATAAACTGATTTTTGCCACCGGC<br>CCGCTGACCGGCACCAGCGCGCCGACCGGTGGCCGCTATATGGTCGTGACCAAATCTCCGCTGA<br>CTGGCTTTATTGCCATGGCGAATAGCGGCGGCTTCTTTGGCGCCGAACTGAAATTTGCGGGCTA<br>CGATGCGATTATTATCGAAGGTAAAGCCGAAAAACCGGTGTACATTTACATTAAAGATGATAAC<br>GTGGAAATTAAAGATGCCTCGCATCTGTGGGGCAAAAAAGTGAGCGAAACCGAAGCGATCCTGC<br>GCGAGGAAATTGGCGATAAAAAAGCGCGCATTGCCAGCATTGGCCCTGCGGGCGAAAATCTGGT<br>GAAGTTTGCCGCAATTATGAATGATGGCCATCGCGCGGCCGGCCGCGGTGGCGTGGGCGCCGTG<br>ATGGGTAGCAAAAACCTGAAAGCGATTGTTGTGCGCGGTTCGAAAAAAGTTGAAGTAGCGGATA<br>AAGAAAAATTCATGCAGGTGGTGCGTGAAAAAATTAACAAACTGCGTAATGATCCTGTTGCAGG<br>CGGCGGCCTGCCGAAATATGGCACCGCGGTGCTGGTGAACATTATTAATGAAAATGGCCTGTAT<br>CCGACACGTAACTTCCAGACCGGCGTATTTGAATATGCGTATGAACAGAGCGGCGAAGCGCTCG<br>CAGCGAAATATCTGGTTCGCAACAAACCGTGCTTTGCCTGCCCGATTGGCTGCGGCCGCGTGAA<br>CAAACTGCCGACTATGGTGGGCGAAACCGAGGGACCGGAATATGAAAGCATTTGGGCGCTGGGC<br>GCCAATTTGGGCATTAACGATCTGGCGAGCATCATTGAAGCCAACCACCTGTGTGACGAATATG<br>GCCTGGACACCATTAGTACCGGTGGCACGCTGGCAGCGGCGATGGAACTGTACGAAAAAGGCCT<br>GATCAAAGATGAAGAACTGGGCGATGGCCCGCCGTTTCGCTTTGGCAATACCGAAGTGCTGCAC<br>TATTATATTGAAAAAATTGCGAAACGTGAAGGCTTTGGCGATAAACTGGCCGAAGGCAGTTATC<br>GCCTGGCGGAATCGTATGGTCACCCGGAATACAGCATGAGCGTGAAAAAACAGGAACTGCCGGC<br>GTATGATCCGCGCGGCGCAGAAGGCCATGGTCTGGAGTATGCCACCAGCAATCGCGGCGGCTGT<br>CACGTAAAAGGCTACATGATTAGCCCGGAAATTCTGGGCTACCCGAAAAAAATGGACCCGCATG<br>ATATTAGCGAAGAAAAAGTGAAAATGGTGAAACTGTTCCAGGATCTGACCGCGCTGATTGATAG<br>CGCGGGTCTGTGTCTGTTCACCAGCTTCGGCCTGGGCGCAGATGATTACCGCGATCTGCTGAAC<br>GCGGCACTGGGCTGGGATTTTAGTACCGAAGAATATCTGAAAATTGGCGAGCGCATTTGGAATG<br>CGGAACGCCTGTTTAACCTGAAAGCGGGCCTGGACCCGGCGAAAGATGATACCTTACCGAAACG<br>TTTTCTGGAAGAACCGATGCCGGAAGGCCCGAACAAAGGCCATGTGGTGCGCCTGAAAGAAATG<br>CTGCCGCGTTATTATAAACTGCGTGGCTGGAACGAAGATGGCAAAATAACCGAAGAAAAACTGA<br>AAGAACTGGGCCTGGAAGAATTCTTATGA |
| *Thermo*AOR3 | ATGTATGGTTATTGGGGTAAAATCCTGCGCGTGAATCTGAGCACCGGCGAAATTAAAGTTGAAG<br>AATTTGATGAGGAACTGGCGAAAAAGTGGCTGGGTAGCCGCGGCCTGGGCATTTACTACCTGCT<br>GAAAGAAATGGACCCGACCGTGGACCCGCTGAGCCCGGAAAATAAATTGATTTTTGCCACCGGC<br>CCGCTCACCGGCACCAGCGCCCCGACCGGCGGGCGTTATATGGTGATTACCAAGAGCCCGCTGA<br>CCGGTTTTATCGCGATGGCGAATAGCGGCGGTTTTTTTGGGGCCGAACTGAAGTTCGCCGGCTA<br>CGATGCCATTATTATTGAAGGCAAAGCGGAAAAGCCGGTTTATATTTATATCAAAGATGATAAC<br>GTGGAAATTCGCGATGCCAGTCACCTGTGGGGCAAAAAAGTCAGCGAAACCGAAGCGATTCTGC<br>GCGAAGAAATTGGCGATAAAAAAGTGCGTATTGCCAGCATCGGCCCGGCGGGCGAAAACCTGGT<br>GAAATTCGCTGCGATTATGAACGATGGCCATCGTGCGGCGGGCCGTGGCGGCGTTGGCGCGGTT<br>ATGGGCAGCAAAAACCTGAAAGCGATTGTTGTGCGCGGCAGCAAGAAAGTTGAAGTTGCAGATA<br>AACAGAAATTTATGAGCGTTGTGCGTGAAAAAATTAACAAACTGCGCAACGATCCGGTTGCGGG<br>TGGCGGTCTGCCGAAATATGGCACCGCCGTGCTGGTAAACATTATTAACGAAAACGGCCTGTAT<br>CCTACCCGTAACTTTCAAACGGGCGTGTTTGAATATGCCTATGAACAATCTGGCGAAGCAATGG<br>CGGCAAAATACCTGGTACGTAACAAACCGTGTTTCGCGTGCCCGATTGGCTGCGGCCGCGTAAA<br>CAAACTGCCGACGGTGGGCGAAACCGAAGGCCCGGAATATGAAAGCATCTGGGCGCTGGGTGCG<br>AACCTGGGTATTAACGATCTGGCCTCTATTATTGAAGCGAACCATCTGTGCGATGAATACGGCT<br>TAGATACCATTTCGACCGGCGGCACTCTGGCGACCGCCATGGAACTGTACGAAAAAGGCCTGAT<br>TAAAGATGAAGAACTGGGCGATGCCCCGCCGTTTCGTTGGGGCAATACCGAAGTGCTGCACTAT<br>TACATCGAAAAAATTGCGAAACGCGAAGGCTTTGGCGACAAACTGGCCGAAGGTGGCTACCGCC<br>TGGCGGAATCGTACGGCCATCCGGAATATTTCATGAGCGTGAAAAAACAGGAACTGCCGGCCTA<br>TGATCCGCGCGGCGCGGAAGGCCATGGCCTGGGCTACGCAACCAACAATCGCGGCGGTTGTCAT<br>ATTAAAAACTACATGATTAGCCCGGAGATTCTGGGCTATCCGTATAAAATGGACCCGCATGATA<br>TTTCGGATGAAAAAGTGAAAATGCTGATTCTGTTTCAGGACCTGACCGCGCTGATTGATAGCGC |

| | |
|---|---|
| | GGGCCTGTGCCTGTTTACGACCTTTGGGCTGGGCGCGGATGATTACCGTGATCTGCTGAATGCG<br>GCCCTGGGCTGGGATTTTAGCACCGAAGATTACTTAAAAATTGGTGAACGCATTTGGAACGCGG<br>AACGTCTGTTTAATCTGAAAGCGGGCCTGGACCCGGCCAAAGATGATACCCTGCCGAAACGCTT<br>TCTGGAGGAACCGATGCCGGAAGGTCCGAACAAAGGCCACGTGGTGCGCTTAAAAGAAATGCTG<br>CCGCGCTATTATGCGCTGCGCGGTTGGACAGAAGATGGTAAAATTCCGGAAGAAAAACTGAAAG<br>AGCTGGGTCTGGAAGAATTTCTGTAA |

**Table S2. Amino acid sequences of proteins used for recombinant expression in *E. coli*.**

| Protein[a] | Amino acid sequence |
|---|---|
| *Pf*AOR | HHHHHHSQDPMYGNWGRFIRVNLSTGDIKVEEYDEELAKKWLGSRGLAIYLLLKEMDPTVDPLS<br>PENKLIIAAGPLTGTSAPTGGRYNVVTKSPLTGFITMANSGGYFGAELKFAGYDAIVVEGKAEK<br>PVYIYIKDEHIEIRDASHIWGKKVSETEATIRKEVGSEKVKIASIGPAGENLVKFAAIMNDGHR<br>AAGRGGVGAVMGSKNLKAIAVEGSKTVPIADKQKFMLVVREKVNKLRNDPVAGGGLPKYGTAVL<br>VNIINENGLYPVKNFQTGVYPYAYEQSGEAMAAKYLVRNKPCYACPIGCGRVNRLPTVGETEGP<br>EYESVWALGANLGINDLASIIEANHMCDELGLDTISTGGTLATAMELYEKGHIKDEELGDAPPF<br>RWGNTEVLHYYIEKIAKREGFGDKLAEGSYRLAESYGHPELSMTVKKLELPAYDPRGAEGHGLG<br>YATNNRGGCHIKNYMISPEILGYPYKMDPHDVSDDKIKMLILFQDLTALIDSAGLCLFTTFGLG<br>ADDYRDLLNAALGWDFTTEDYLKIGERIWNAERLFNLKAGLDPARDDTLPKRFLEEPMPEGPNK<br>GHTVRLKEMLPRYYKLRGWTEDGKIPKEKLEELGIAEFY |
| *Thermo*AOR5 | HHHHHHSQDPMMGYMGKILRVNLTTGEIKVEELDEELAKKFIGGRGLASKYLYDEIDPTVDPLS<br>PENKLIFATGPLTGTTAPTGGRYMVVTKSPLTGFIASSNSGGFFGPELKFAGYDMLIVEGKADK<br>PVYISIKDDNVEIKDASHLWGKKVSETTDILLEEFGDKKARVACIGPAGENLVKFAAIMNDKHR<br>AAGRSGVGAVMGSKNLKAIVVRGSKKVEVADKEKFMEVVKEKIKKIKENPVTGGGLPKYGTAVL<br>VNIINENGLYPTRNFQTGVFEYAYEQSGEALAAKYLVRNKPCFACPIGCGRVSKLPAVGETEGP<br>EYESIWALGANCGINDLASIVEANHLCDEYGLDTISTGGTLAAAMELYEKGIIKEEELGDGPPL<br>RFGNSEVLHYYIEKIAKREGFGDKLAEGSYRLAESYGHPEYSMSVKKQELPAYDPRGAEGHGLE<br>YATSNRGGCHVRGYMISPEILGVPKKLDPHDISEEKAKWVKIFQDLTAVIDSAGLCLFTSFALG<br>ADDYRDLLNAATGFDYTTEEMMKAGERIWNLERLFNLKAGLDPEKDDTLPKRFLEEPMPEGPNK<br>GHVVRLKELLPKYYKLRGWNEDGIPTEEKLKELGLAEFY |
| *Thermo*AOR4 | HHHHHHSQDPMMGYMGKILRVNLTTGEIKVEELDEELAKKWIGGRGLGTYYLLKEMDPKVDPLS<br>PENKLIFATGPLTGTSAPTGGRYMVVTKSPLTGFIAMANSGGFFGAELKFAGYDAIIIEGKAEK<br>PVYIYIKDDNVEIKDASHLWGKKVSETEAILREEIGDKKARIASIGPAGENLVKFAAIMNDGHR<br>AAGRGGVGAVMGSKNLKAIVVRGSKKVEVADKEKFMQVVREKINKLRNDPVAGGGLPKYGTAVL<br>VNIINENGLYPTRNFQTGVFEYAYEQSGEALAAKYLVRNKPCFACPIGCGRVNKLPTMVGETEG<br>PEYESIWALGANLGINDLASIIEANHLCDEYGLDTISTGGTLAAAMELYEKGLIKDEELGDGPP<br>FRFGNTEVLHYYIEKIAKREGFGDKLAEGSYRLAESYGHPEYSMSVKKQELPAYDPRGAEGHGL<br>EYATSNRGGCHVKGYMISPEILGYPKKMDPHDISEEKVKMVKLFQDLTALIDSAGLCLFTSFGL<br>GADDYRDLLNAALGWDFSTEEYLKIGERIWNAERLFNLKAGLDPAKDDTLPKRFLEEPMPEGPN<br>KGHVVRLKEMLPRYYKLRGWNEDGKITEEKLKELGLEEFL |
| *Thermo*AOR3 | HHHHHHSQDPMYGYWGKILRVNLSTGEIKVEEFDEELAKKWLGSRGLGIYYLLKEMDPTVDPLS<br>PENKLIFATGPLTGTSAPTGGRYMVITKSPLTGFIAMANSGGFFGAELKFAGYDAIIIEGKAEK<br>PVYIYIKDDNVEIRDASHLWGKKVSETEAILREEIGDKKVRIASIGPAGENLVKFAAIMNDGHR<br>AAGRGGVGAVMGSKNLKAIVVRGSKKVEVADKQKFMSVVREKINKLRNDPVAGGGLPKYGTAVL<br>VNIINENGLYPTRNFQTGVFEYAYEQSGEAMAAKYLVRNKPCFACPIGCGRVNKLPTVGETEGP<br>EYESIWALGANLGINDLASIIEANHLCDEYGLDTISTGGTLATAMELYEKGLIKDEELGDAPPF<br>RWGNTEVLHYYIEKIAKREGFGDKLAEGGYRLAESYGHPEYFMSVKKQELPAYDPRGAEGHGLG<br>YATNNRGGCHIKNYMISPEILGYPYKMDPHDISDEKVKMLILFQDLTALIDSAGLCLFTTFGLG<br>ADDYRDLLNAALGWDFSTEDYLKIGERIWNAERLFNLKAGLDPAKDDTLPKRFLEEPMPEGPNK<br>GHVVRLKEMLPRYYALRGWTEDGKIPEEKLKELGLEEFL |

[a] The $His_6$ tag is shown in green and the linker in orange.

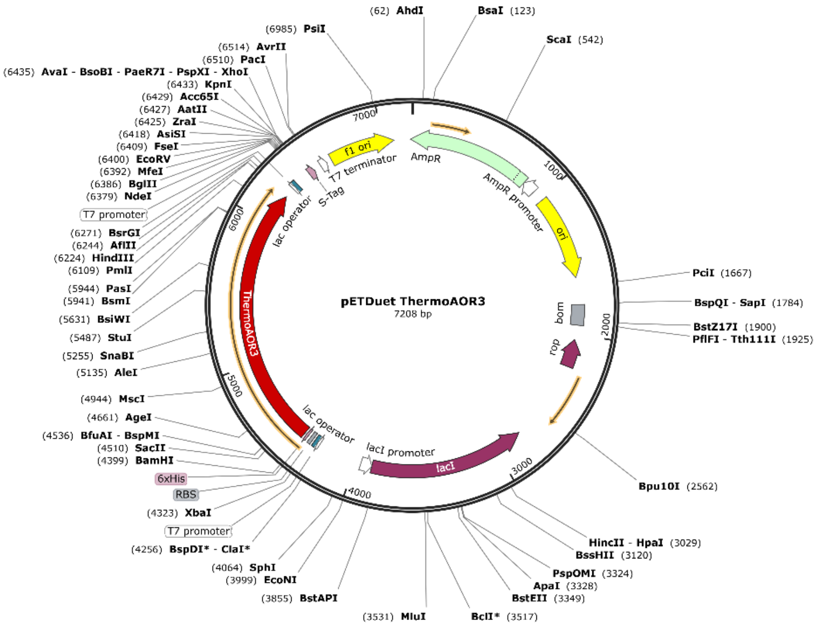

**Figure S2. Plasmid map of pETDuet-1 *Thermo*AOR3.**

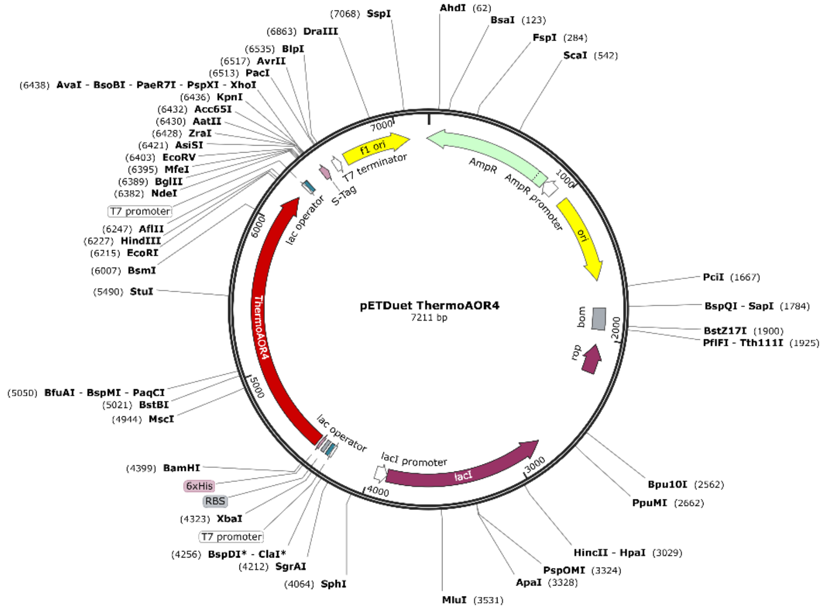

**Figure S3. Plasmid map of pETDuet-1 *Thermo*AOR4.**

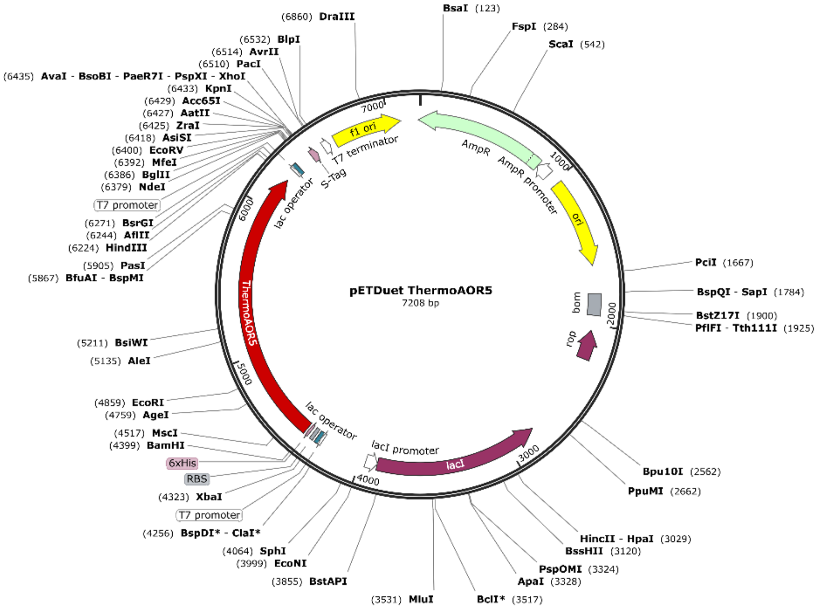

**Figure S4. Plasmid map of pETDuet-1 *Thermo*AOR5.**

**Table S3. Aldehyde substrates used for the substrate screening of *Thermo*AOR3.**

| Substrate type | Substrate[a] | Solvent | Supplier |
|---|---|---|---|
| Linear aliphatic | DL-Glyceraldehyde-3-phosphate | Water | Merck – Sigma-Aldrich |
| | Acetaldehyde | Water | Merck – Sigma-Aldrich |
| | Butyraldehyde | Water | Merck |
| | Pentanal | Water | Fluka Chemicals |
| | Hexanal | DMSO | Merck – Sigma-Aldrich |
| | Octanal | DMSO | TCI Europe N.V. |
| | Decanal | DMSO | Merck – Sigma-Aldrich |
| | Dodecanal | DMSO | Merck – Sigma-Aldrich |
| Branched chain aliphatic | Isobutyraldehyde | Water | Acros Organics |
| | 2-methylbutyraldehyde | DMSO | Merck – Sigma-Aldrich |
| | 3-3-dimethylbutyraldehyde | Water | Merck – Sigma-Aldrich |
| | 2-methylvaleraldehyde | DMSO | Merck – Sigma-Aldrich |
| | Isovaleraldehyde | Water | Merck – Sigma-Aldrich |
| | 2-ethylbutyraldehyde | DMSO | Merck – Sigma-Aldrich |
| | 2-ethylhexanal | DMSO | Merck – Sigma-Aldrich |
| | Citral | DMSO | Alfa Aesar |
| Aromatic | Vanillin | Water | Acros Organics |
| | Hydrocinnamaldehyde | DMSO | Merck – Sigma-Aldrich |
| | 2-Phenylpropionaldehyde | DMSO | Merck – Sigma-Aldrich |
| | 4-nitrobenzaldehyde | DMSO | Fluka Chemicals |
| | benzaldehyde | DMSO | Merck – Sigma-Aldrich |
| Other | Cis-9-octadecenal | DMSO | Merck – Sigma-Aldrich |
| | Furfural | Water | Merck – Sigma-Aldrich |
| | 5-hydroxymethylfurfural | Water | Acros Organics |
| Reference | Crotonaldehyde | Water | Avantor |

[a] Substrates which were not soluble in water, were dissolved in dimethylsulfoxide (DMSO). All stock solutions were diluted 80 times in the reaction, to a final concentration of 0.5 mM. Therefore, the final concentration of DMSO, if necessary, in the reaction was 1.25% (v/v).

**Table S4. GC conditions used for crotonaldehyde and crotonic acid measurements.**

| Column | Temperature profile | Retention time (min) |
|---|---|---|
| GC | 95°C, hold 3.0 min | Crotonaldehyde: 4.46 |
| CP-Wax 52 CB (Agilent) | 10°C/min to 105°C, hold 1 min | n-Dodecane (IS): 6.30 |
| (25 m, 0.25 mm, 1.2 μM) | 30°C/min to 207°C, hold 1 min | DMSO: 10.15 |
| Carrier gas $N_2$[a] | 20°C/min to 230°C, hold 1 min | Crotonic acid: 11.08 |
| | 30°C/min to 245°C, hold 2 min | |

[a]Linear velocity used is 30 cm/sec.

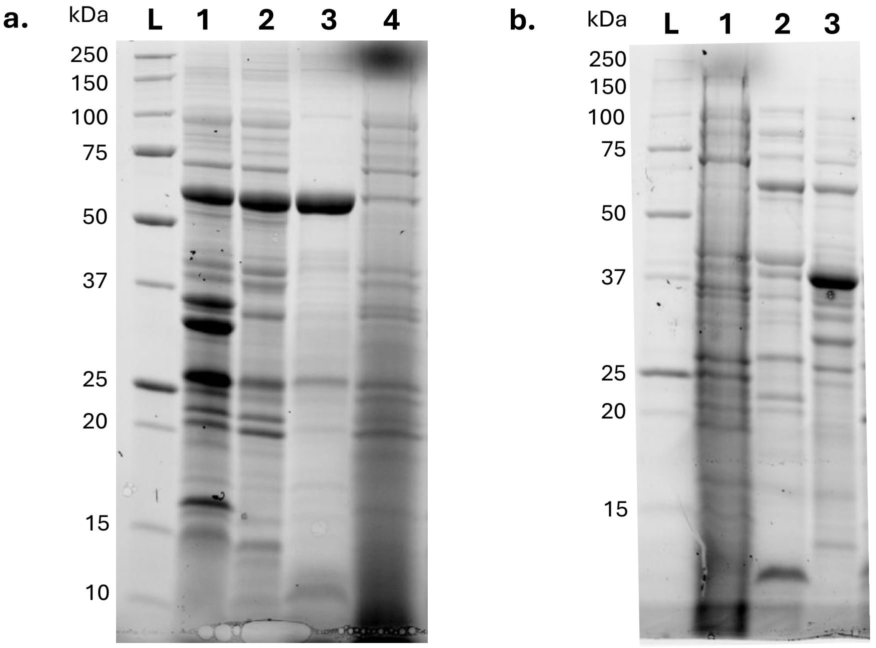

**Figure S5. Analysis of recombinant expression of *Pf*AOR in different *E. coli* strains.**

SDS-PAGE of heat purified *P. furiosus* AOR recombinantly expressed in *E. coli* Rosetta-Gami 2(DE3, **a**) and MC1061 (**b**). L. Precision Plus Protein Standard Unstained (Bio-Rad Laboraties Inc., Hercules, CA, USA). **a**. 1. 30 μg cell pellet. 2. 30 μg cell-free extract before purification. 3. 5 μg heat purified *Pf*AOR. 4. Rosetta-Gami 2(DE3) cells before induction. **b**. 1. MC1061 cells before induction. 2. 30 μg cell-free extract. 3. 30 μg cell pellet.

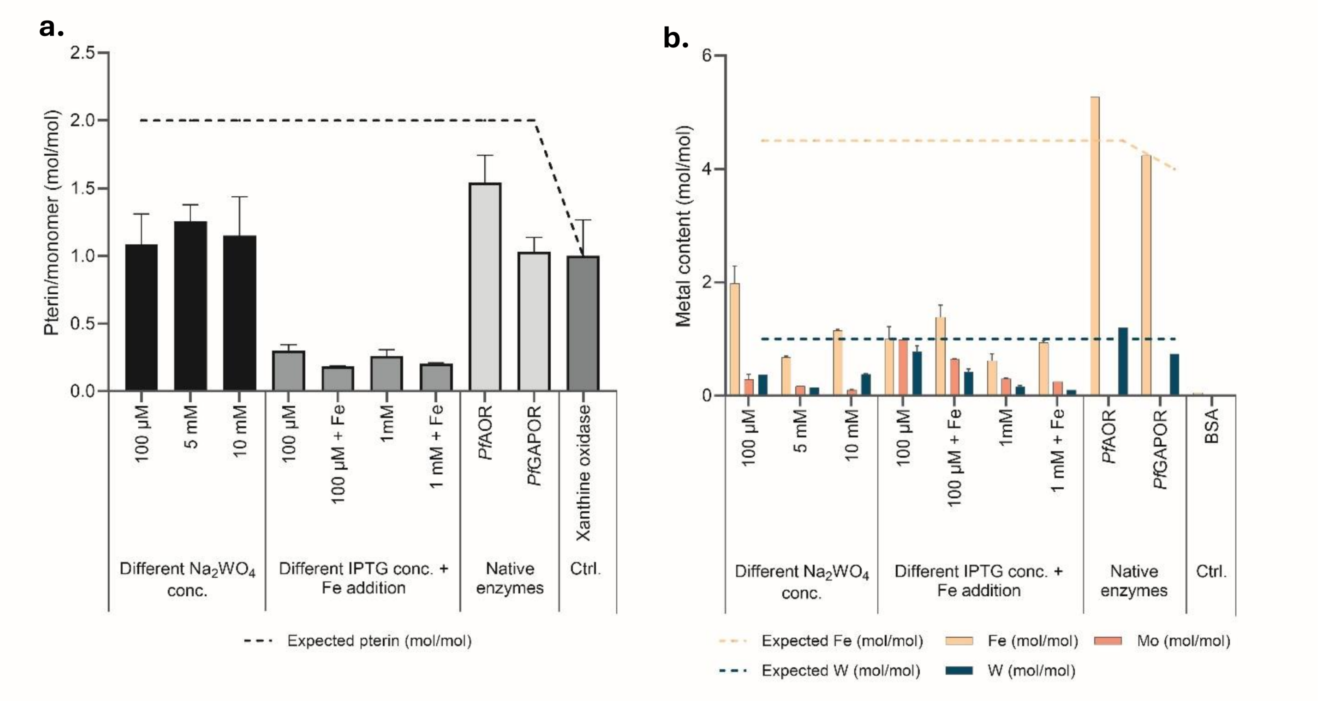

**Figure S6. Pterin and metal content of *Pf*AOR proteins.**

Pterin and metal content present in the W-bis-MPT cofactor and [4Fe-4S]-cluster were measured upon the addition of different concentrations of medium additives ($Na_2WO_4$, IPTG, and Fe). The recombinantly produced proteins are compared to AOR and GAPOR purified from *P. furiosus* cells. **a**. Fluorescence assay with a $\lambda_{ex}$ of 360 nm and a $\lambda_{em}$ of 460 nm. The pterin content measured in different assays is normalized to a pterin content of Xanthine oxidase of 1 mol/monomer. **b**. ICP-MS analysis to measure Molybdenum (Mo), Iron (Fe), and tungsten (W). Bovine serum albumin (BSA) was used as a negative control.

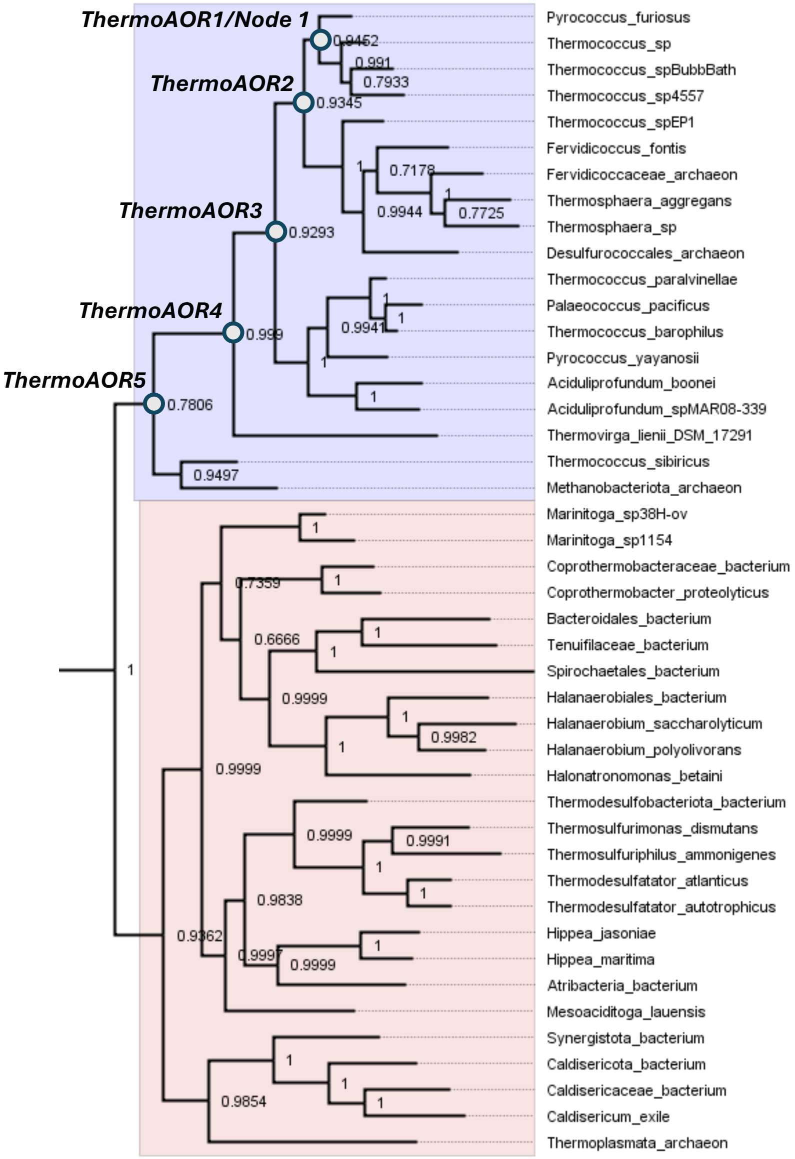

**Figure S7. Phylogenetic tree of *Pf*AOR used for ancestral sequence reconstruction.**

The wild-type homologue enzymes used for phylogenetic analysis are labeled according to the organism they originate from. The selected nodes of *Pf*AOR have been labeled with blue circles and their posterior probabilities are shown. The proteins are numbered and named according to the distance from the *Pf*AOR, ranging from *Thermo*AOR1 (Node 1, closest to *Pf*AOR) to *Thermo*AOR5 (Node 5, furthest from *Pf*AOR). The archaeal part of the tree is highlighted in light purple, while the bacterial part is depicted in light orange. *Pf*AOR, the reference protein, is at the top of the phylogenetic tree.

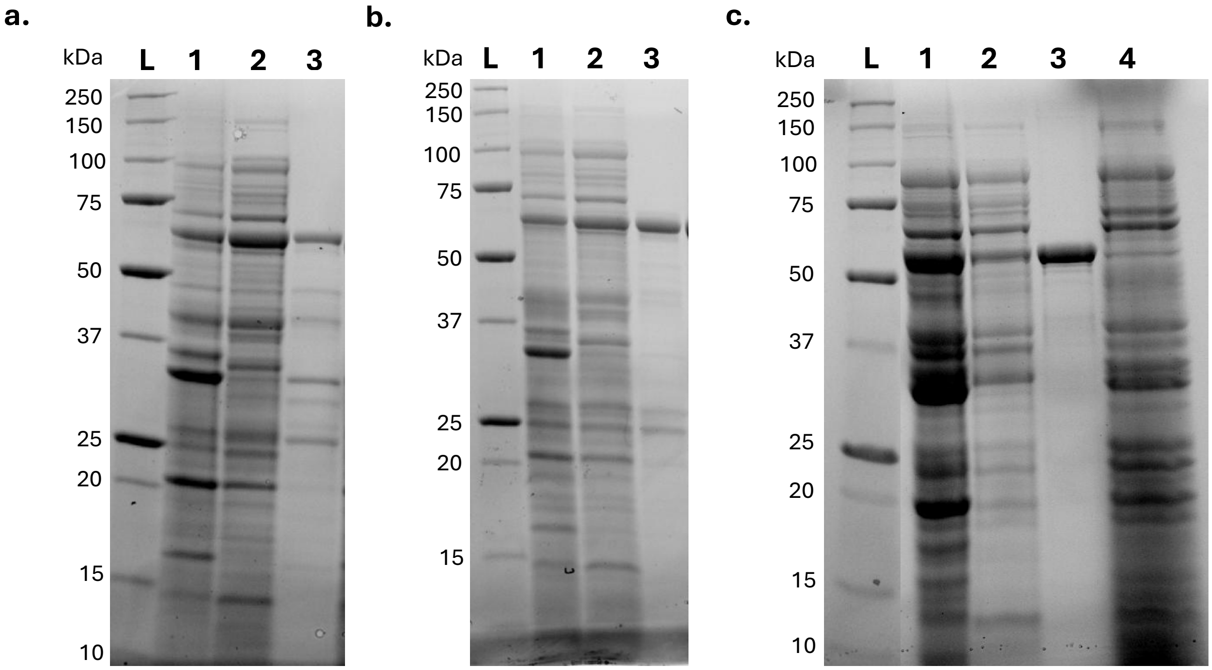

**Figure S8. SDS-PAGE analysis of the recombinant expression and purification of *Thermo*AOR5 (a), *Thermo*AOR4 (b) and *Thermo*AOR3 (c).**

All proteins have been separated and visualized by SDS-PAGE analysis. L. Precision Plus Protein Standard Unstained (Bio-Rad Laboraties Inc., Hercules, CA, USA).1. 30 µg cell pellet. 2. 30 µg cell-free extract. 3. 5 µg purified protein. 4. *E. coli* Origami 2(DE3) cells before induction.

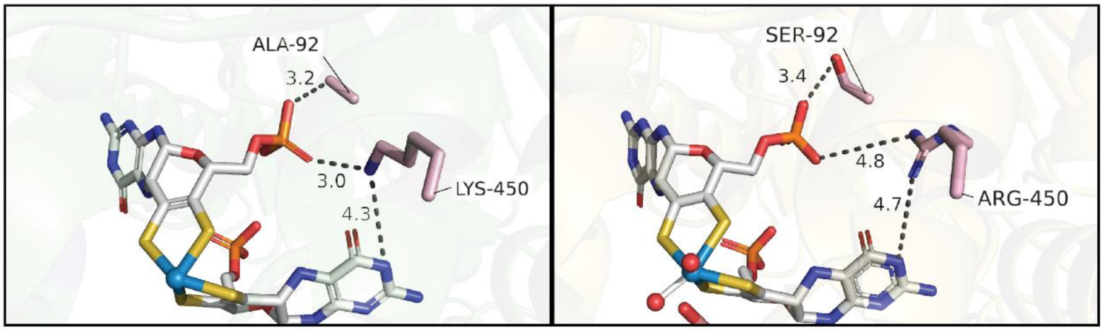

*Pf*AOR *Thermo*AOR5

**Figure S9. Active site mutations present in *Thermo*AOR5.**

Only in *Thermo*AOR5 two amino acids close to the pterin cofactor are mutated. Ala92 and Lys450 in *Pf*AOR are mutated to Ser92 and Arg450 in *Thermo*AOR5.

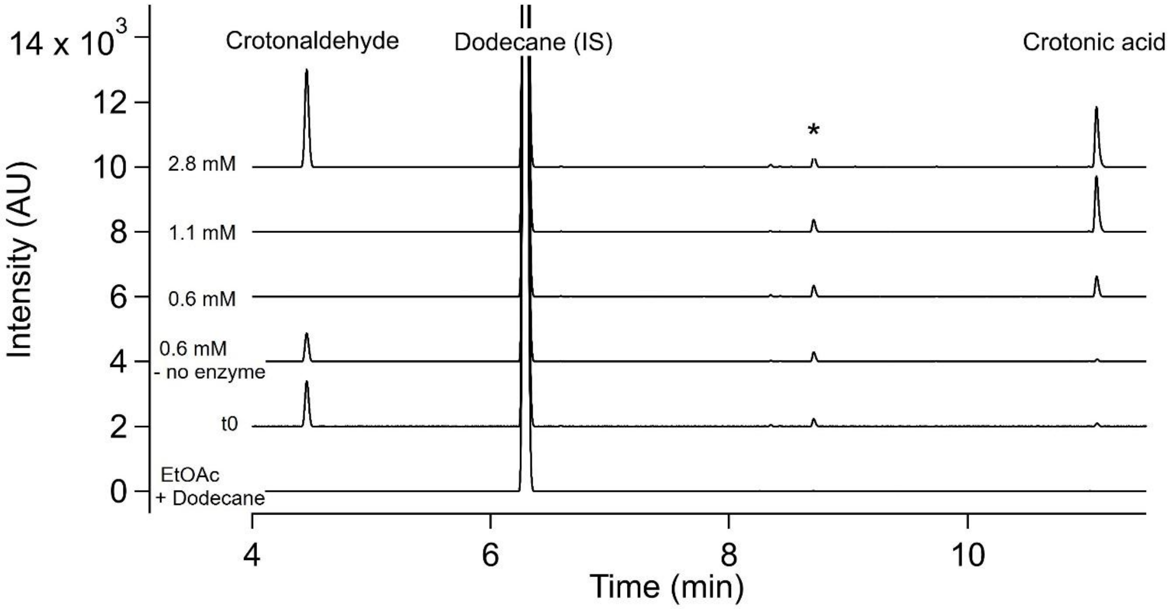

**Figure S10. Gas chromatograms of AOR catalyzed crotonaldehyde oxidation.**

Retention times; Crotonaldehyde: 4.46. n-Dodecane (IS): 6.30. DMSO: 10.15. Crotonic acid: 11.08. * Substrate impurity.

### *Thermo*AOR3 temperature profile using sonication as cell lysis method

The proteins used in this manuscript were lysed using a chemical lysis method with FastBreak™ lysis reagent. When changing this lysis method to lysis using sonication, the activity of purified *Thermo*AOR3 increased approximately 2-fold (0.8 U/mg vs 2 U/mg at 60°C), as well as the stability of the enzyme at temperatures above 70°C (Fig. 6).

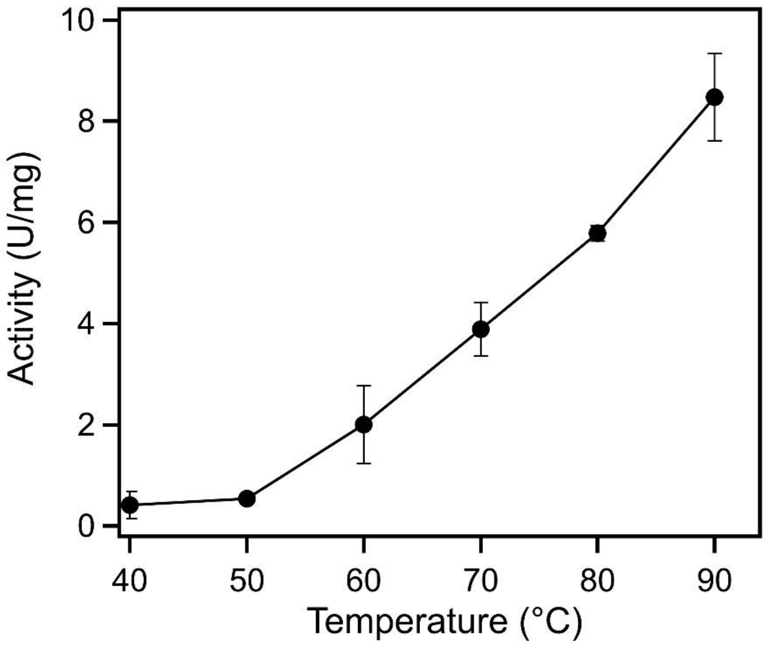

**Figure S11. Initial activity screening of *Thermo*AOR3 using different temperatures.**

Origami 2(DE3) cells were lysed using sonication after which *Thermo*AOR3 was purified and used for activity screening using BV as electron acceptor. The assay contained 3 mM BV in EPPS buffer (50 mM, pH 8.4), 0.2 µM purified *Thermo*AOR3, and 1 mM crotonaldehyde. Activity was calculated based on the absorbance increasement at 600 nm using ε($BV^{+\cdot}$) =7,400 $M^{-1}cm^{-1}$.

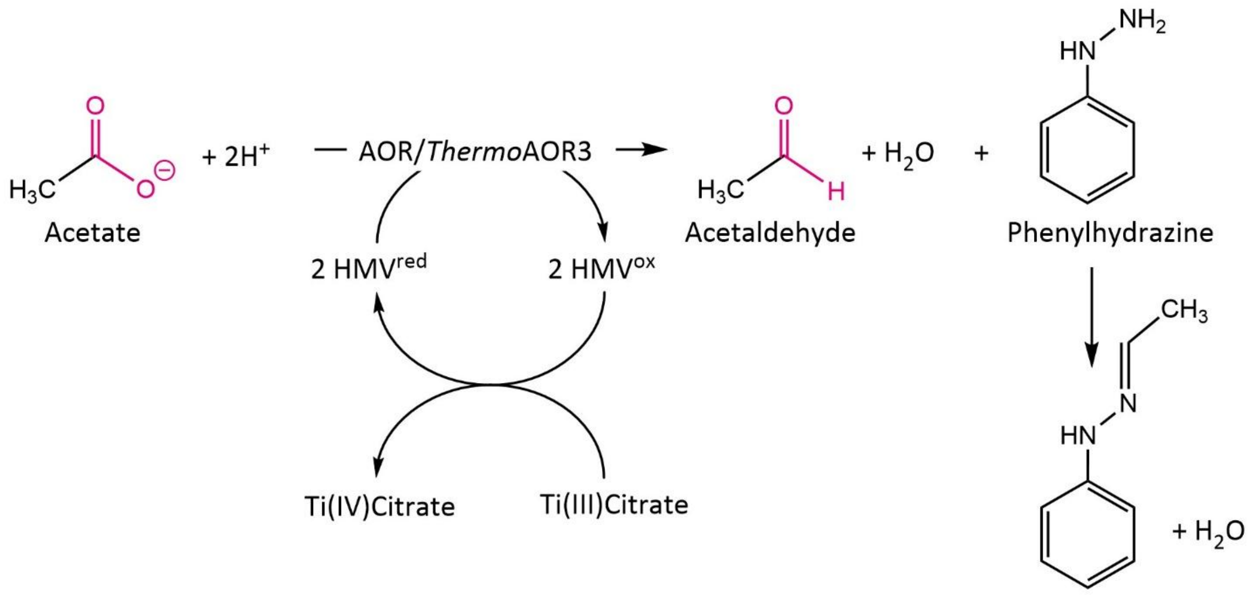

**Figure S12. Carboxylic acid reduction assay using *Thermo*AOR3.**

Acetate is reduced by *Thermo*AOR3 using hexamethylviologen (HMV) as electron donor. HMV is pre-reduced using Ti(III)Citrate. A favorable equilibrium is created by trapping the produced acetaldehyde by binding to phenylhydrazine. HMV – Hexamethylviologen. Red – reduced. Ox- oxidized.
